# Assessing Hardy-Weinberg Equilibrium in T2T-aligned 1000 Genomes Project

**DOI:** 10.64898/2026.01.05.696401

**Authors:** Elika Garg, Jaffa Romain, Lei Sun, Andrew D. Paterson

## Abstract

Quality control of markers in genome-wide association studies often includes testing for Hardy-Weinberg equilibrium (HWE). However, this is usually implemented in a homogeneous population without stratifying by sex. Previous work indicates sex-based selection at numerous autosomal loci in cohorts with active recruitment. Sex chromosome sequences can also interfere with autosomal SNPs. We examined genome-wide sex-specific HWE deviations across populations in the telomere-to-telomere (T2Tv2)-aligned high-coverage whole genome sequence of the 1000 Genomes Project data of 2,490 individuals. Our analysis was restricted to bi-allelic SNPs with non-missing genotypes and MAF>=5% in both sexes of the five super-populations. We employed a robust allele-based approach for HWE testing, which enabled the quantification of directional deviations from HWE. A second-order omnibus meta-analysis combining results from the five super-populations and both sexes revealed that 0.9% autosomal SNPs exhibited a significant deviation from HWE at p<5e-8. Most of these deviations were found to be associated with genomic features relating to poor sequence quality. Filtering results to reliable genomic regions yielded 255 autosomal and 1 NPR X chromosomal SNPs, of which 140 autosomal SNPs also showed significant heterogeneity across populations but not across sexes. 8 SNPs in a 15-bp region on chr14 showed excess heterozygosity in both sexes of the AFR (African) super-population. We also generated a well-performing multivariate predictor of HWD (deviation from HWE) using multiple sequence features, which could be combined with HWD estimates in future studies to select SNPs that deviate from HWE due to technical rather than biological reasons.

**Author Summary:** We conducted a specific quality control test, which compares the observed and expected genotype counts, on an updated version of the 1000 Genomes Project whole genome sequence data generated on ∼2500 individuals. We first performed this analysis by grouping the data by ancestry and sex. We then combined and contrasted the group results. We found that most regions that differed between observed and expected counts overlapped regions of the genome which are difficult to sequence using current short read technology. In the remaining regions we found an interesting cluster of SNPs in a single ancestry, where there is a gross excess of heterozygous genotypes. GWASes typically use a standard strict threshold for this quality control test for genotyping arrays to remove SNPs. Here we suggest a more nuanced approach that is applicable to whole genome sequence data.

## Introduction

### Sex-specific alignment of 1000 Genomes Project (1KGP) data using the complete T2T reference genome

Data from the 1000 Genomes Project has been widely used for genetics research (1). Over the years, 1KGP has generated numerous versions, including data from genome-wide association studies (GWAS) arrays and exome sequencing. A recent release of 1KGP consists of 3,202 high-coverage short-read whole genome sequenced samples aligned to GRCh38 (GCA_000001405.15; (2)). This data was then first re-aligned to the Telomere-to-Telomere reference genome (T2T-CHM13v1.0; (3,4), which did not include chromosome Y (chrY)). Data aligned to T2T was shown to improve autosomal mappability and variant calling compared to GRCh38. The same high-coverage 1KGP data has since been realigned to T2T-CHM13v2.0, which includes chrY (5); we refer to this version as “v2”.

Sex chromosomes, X and Y, have a common evolutionary origin and consequently share regions of high sequence similarity (5), including, but not limited to, the pseudo-autosomal regions (PARs). Therefore, aligning short-read sequencing data to a reference genome in a sex-chromosome agnostic manner can lead to misalignments. v2 aligns XX and XY samples to sex-chromosome-aware references using XYalign (6). To reduce misalignments, XYalign masks chrY of the reference genome for XX samples, and chrY PARs for XY samples (5). However, extensive quality assessment of v2 data, specifically the Hardy-Weinberg equilibrium (HWE), has not been reported.

### Hardy-Weinberg equilibrium (HWE)

HWE is a fundamental principle in population genetics that quantifies the relationship between allele and genotype frequencies (7,8). A SNP is in HWE if the genotype frequencies depend only on the allele frequencies in a given population. Deviation from HWE can be due to either technical reasons or the violation of HWE assumptions, such as mutation, migration, non-random mating, or selection (9). HWE is particularly important in the quality control of GWAS, where deviation from HWE can raise concerns about data quality, selection, or underlying population structure. However, not all deviation is artefactual: it can be driven by disease association (10,11), participation bias (12) or admixture (13). Graffelman et al., 2017 performed a genome-wide study of HWE in the Japanese and Yoruban populations from the phase 3 1KGP, which was aligned to GRCh37, and they reported more HWE deviation than would be expected by chance alone. They also indicated there is a complex relationship between read depth and deviation from HWE, where most of the SNPs were derived from low-coverage WGS. Apart from that, there has been surprisingly little emphasis on HWE in 1KGP, despite the use of this data for many purposes which often rely upon the assumption of HWE (1,2).

### Genotyping errors

Systematic sequencing and alignment biases can lead to genotyping errors and interfere with true biological signals in genetic data (14). Common errors in short-read sequencing arise when reads align to multiple regions of the genome with high sequence similarity and result in incorrect SNP calls (15). For example, autosomal sequences can misalign to chrY in males, leading to sex differences in genotyping errors due to a paralogous sequence variant (PSV; (16,17)). Such misalignment can also occur due to segmental duplications, low-complexity regions and repeats. The Genome in a Bottle consortium (GIAB) has categorized certain genomic regions as “difficult-to-sequence” regions, where confidence in calls is low (18). In contrast, Illumina has defined short-read accessibility masks to mark regions where SNP discovery and genotyping are relatively reliable (19). Ogata et al., 2023 have comprehensively assembled other lists of problematic sequence features which, if excluded, can improve the accuracy of biological signals (20).

### Sex differences in minor allele frequency (sdMAF)

Recent studies have reported sdMAF of SNPs on but not limited to chromosome X (chrX; (12,17,21)). sdMAF on chrX in PARs and at PAR-NPR boundaries are likely due to sex-linkage (17,22). But sdMAF can also be driven by genotyping errors, for example, PSVs causing cross-alignment of genomic regions between autosomes and sex chromosomes (17).

Pirastu et al., 2021 performed an autosomal GWAS for sex using data from 2.46 M customers of 23andMe, having previously shown that sex was an autosomally heritable trait in studies with active as opposed to passive participation. They identified 158 autosomal loci associated with sex and examined them further. Included in the follow-up analyses was HWE testing (p<10e-6), but this was only performed in sex-combined samples, despite the observed sdMAF. Combining groups with different MAFs can lead to Hardy–Weinberg disequilibrium.

### HWE in recently-admixed populations

In recently-admixed populations, it has been theoretically demonstrated that achieving HWE on chrX takes more than one generation, unlike the autosomes (13,23,24). However, this has never been demonstrated in practice. Since 1KGP has some recently-admixed populations (e.g. African Ancestry in Southwest US [ASW], African Caribbean in Barbados [ACB]), we compare HWD (deviation from HWE) in these populations between autosomes and NPR.

In conclusion, although HWE evaluation has been extensively used for quality control in large-scale genotyping/sequencing studies, it has not been previously studied in the high-coverage short-read whole genome sequences of 1KGP aligned to T2T-CHM13v2.0. Here, we perform sex- and super-population-stratified HWD analyses in this data, and we then combine the results using meta-analysis. SNPs identified as deviating from HWE were then subjected to bioinformatic annotations. Additionally, we used these annotations to generate a multivariate predictor of HWD.

## Results

### 1000 Genomes Project data

The high coverage 1KGP data which has been re-aligned in a sex-specific manner to the T2T-CHM13v2.0, referred to here as “v2”, consists of 2,490 unrelated samples (1,263 females and 1,227 males) of which a total of 11 million bi-allelic non-missing SNPs were first selected irrespective of MAF (10,984,307 in autosomes, 383,435 in NPR, 16,217 in PAR1, and 1,006 in PAR2). Table 1 shows the number of males and females in each of the five super-populations (ranging from 170 males in AMR to 336 females in AFR), along with the number of SNPs with MAF>=5% in each group (ranging from 4,987,954 in EAS males to 7,918,101 in AFR females).

**Table 1.**
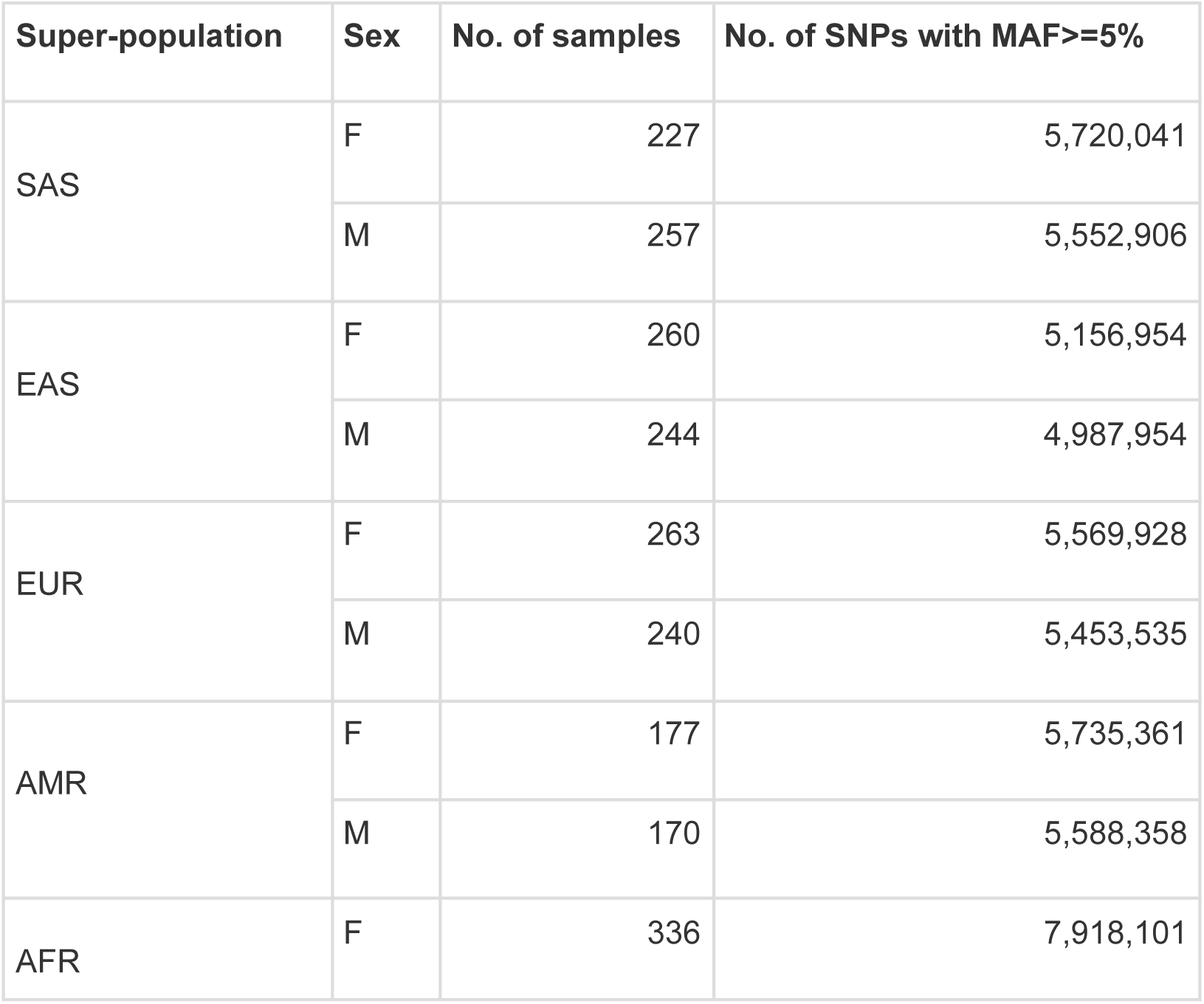
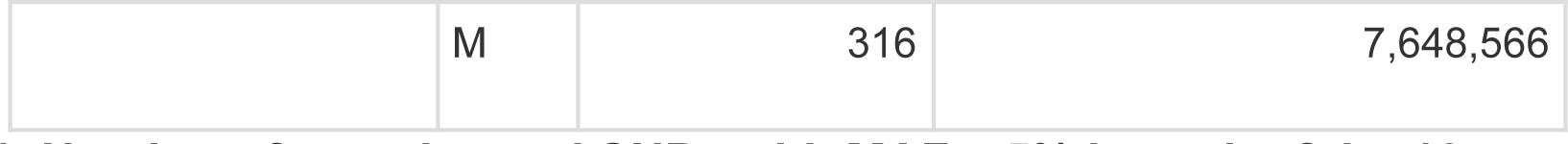
Number of samples and SNPs with MAF > 5% in each of the 10 groups. Groups are made on the basis of super-population (SAS = South Asian, EAS = East Asian, EUR = European, AMR = Admixed American, AFR = African) and sex (F = females, M = males). Note that SNP counts in females include NPR, which is absent in males.

### HWD stratified by sex and super-population and meta-analyses

Genome-wide significant deviations from HWE (HWD) as measured by p<5e-8, were found across the genome in all five sex-stratified super-populations. Figure 1 shows the HWE p-values in each of the 10 groups (5 super-populations x two sexes) in autosomal SNPs with MAF>=5% in each group (see Table 1). HWD in the autosomes had similar patterns across groups, for example, at telomeres and around centromeres.

**Figure 1.**
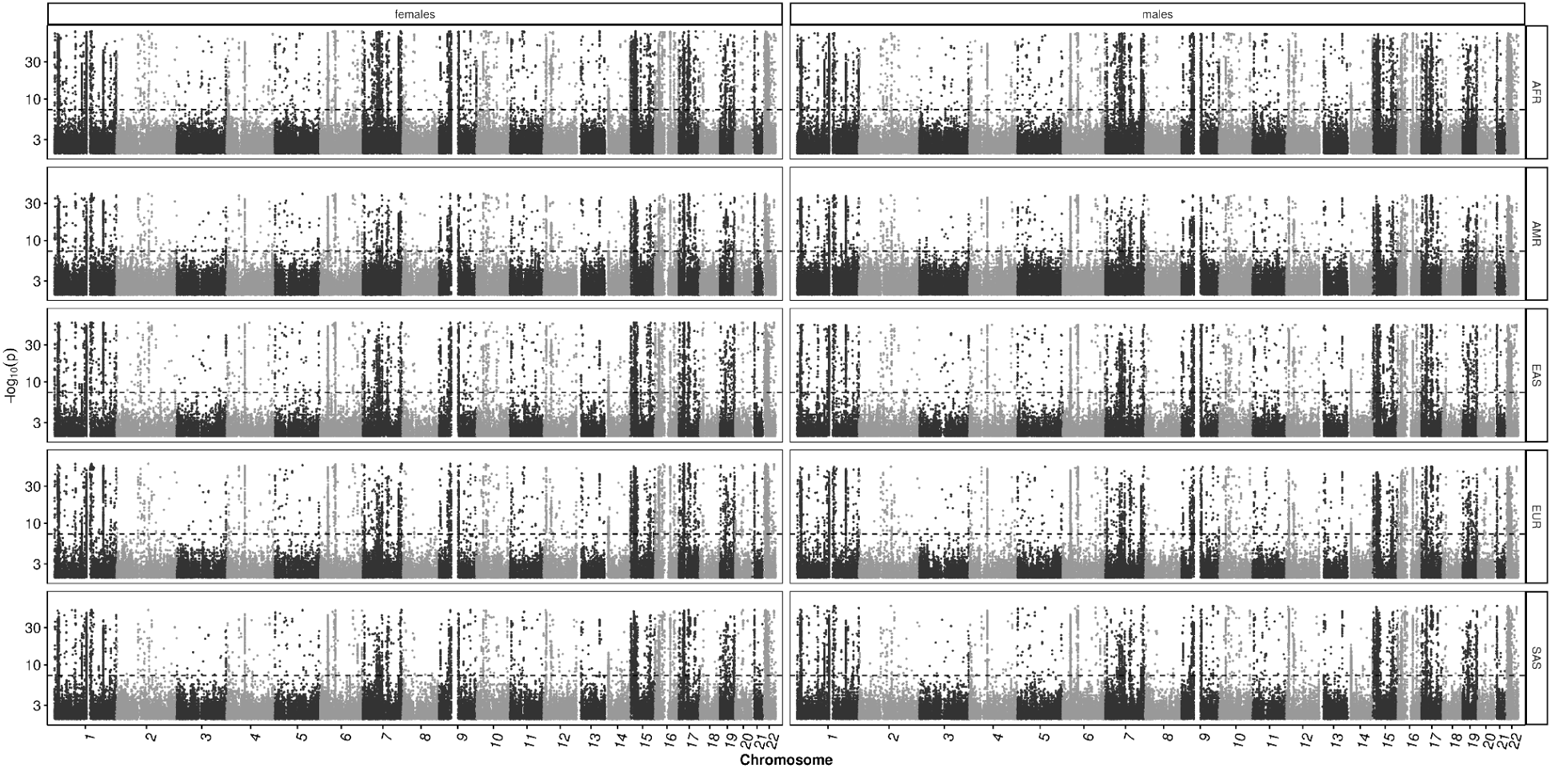
Manhattan plots for autosome-wide HWE in each of the 10 groups. HWE results from five super-populations and two sexes are shown for autosomal SNPs with MAF>=5%. Dashed line represents the genome-wide significant threshold of p=5e-8. Y-axis is presented on a log10 for SNPs with HWE p-value<0.01.

Figure 2a shows the distribution of HWE p-values from the omnibus meta-analysis of all 10 groups for 3,377,201 autosomal SNPs (0.9% SNPs are significant, Table 2) that are in common at MAF>=5% in each group. Figure 2b shows the HWE heterogeneity results for the same set of SNPs. Overall, as expected, there was more consistency between sexes of the same super-population than between super-populations of the same sex. Table 2 presents the overlap between SNPs identified in the HWE omnibus and heterogeneity meta-analyses, stratified by the significance of each test. About half of the significant SNPs in the omnibus meta-analysis also showed significant HWE heterogeneity. Interestingly, seven SNPs exhibited significant HWE heterogeneity but were not significant in the omnibus test, as expected when directional effects differ across groups. These SNPs mapped to two loci on chromosomes 2 and 4 (Tables 2 and S1). At the region on chr2, the most pronounced deviation from HWE was observed in AMR females, characterized by a paucity of heterozygotes, and the HWE heterogeneity was driven by super-populations within females. In contrast, in the region on chr4, the most pronounced deviation from HWE was observed in AFR males, characterized by an excess of heterozygosity (Table S1).

**Figure 2.**
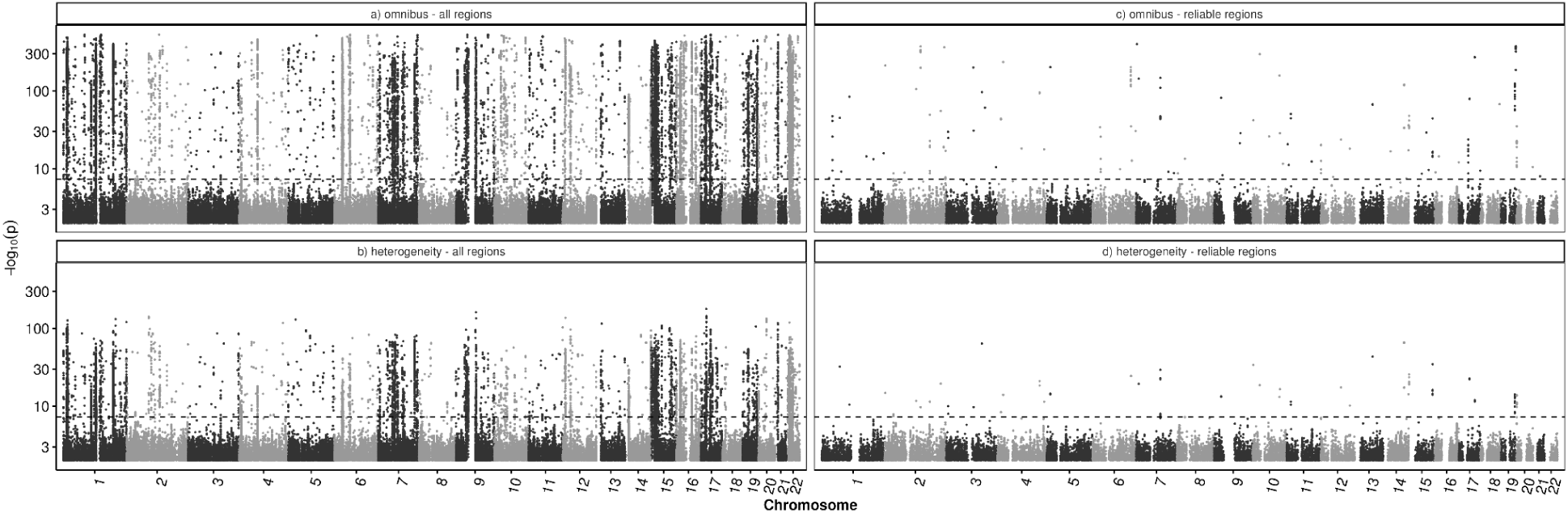
Manhattan plots for autosome-wide HWE meta-analysis of 10 groups. Meta-analysis results from five super-populations and two sexes are shown for autosomal SNPs which have MAF>=5% in all groups: a) omnibus and b) HWE heterogeneity results for all regions (n=3,377,201), c) omnibus and d) HWE heterogeneity results for reliable regions (n=1,347,995). Dashed line represents the genome-wide significant threshold of p=5e-8. Y-axis is presented on a log10 scale for SNPs with meta-analysis p-value<0.01.

**Table 2.**
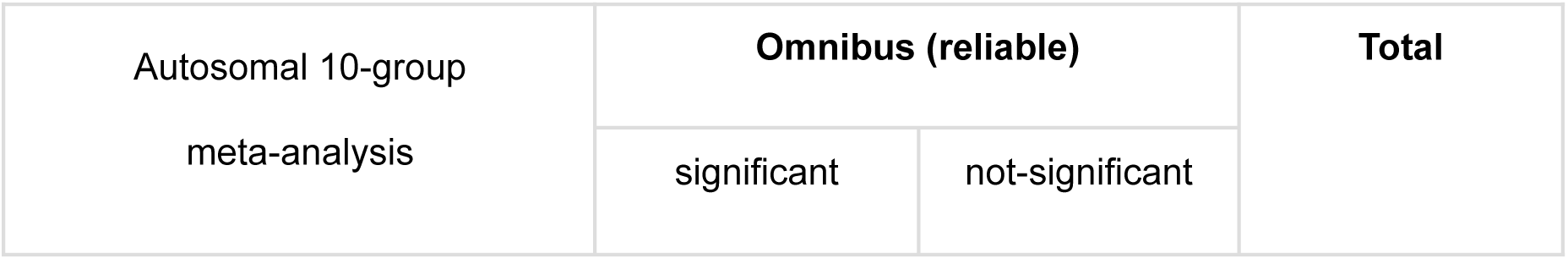
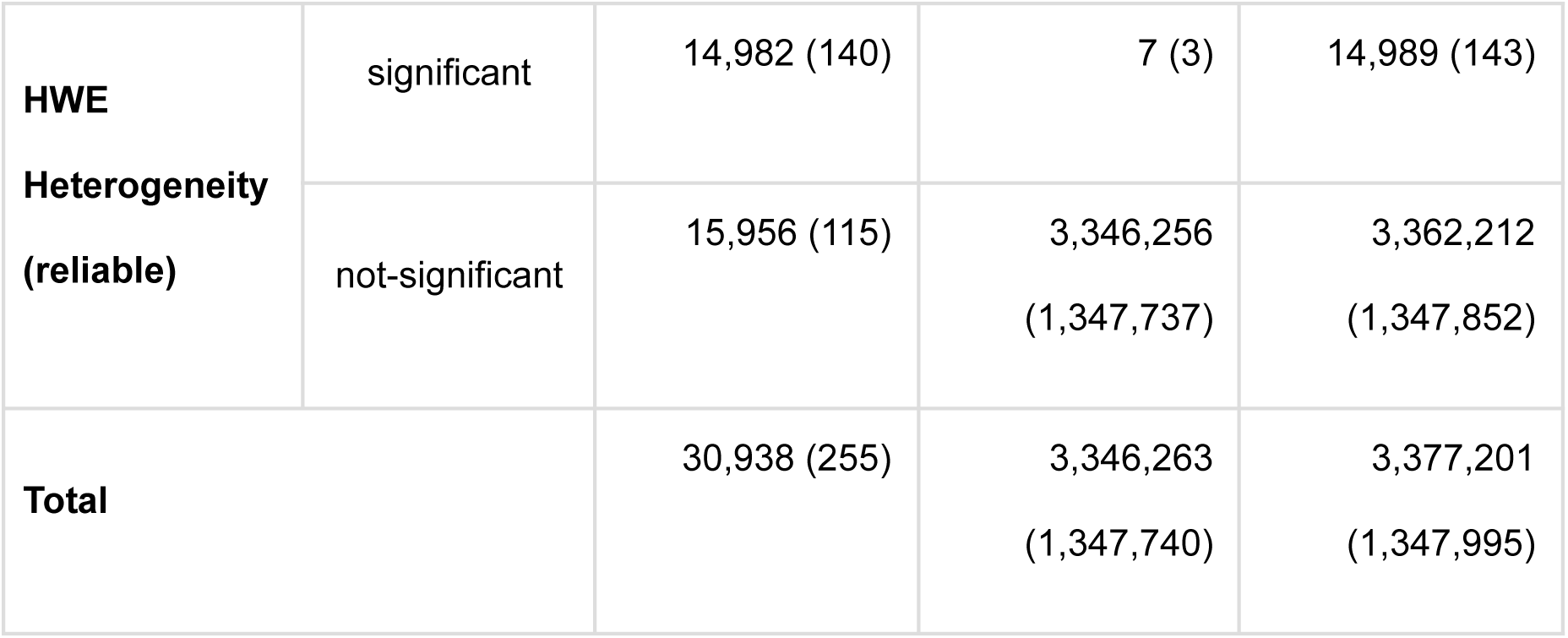
Overlap of autosome-wide SNPs between HWE meta-analyses of 10 groups. Meta-analyses results from five super-populations and two sexes are compared for autosomal SNPs which have MAF>=5% in all groups. The overlap of significant and not-significant SNPs between HWE heterogeneity and omnibus meta-analyses are shown. Numbers in brackets indicate the subset of SNPs in reliable regions.

| Autosomal 10-group<br>meta-analysis | Omnibus (reliable) |  | Total |
| --- | --- | --- | --- |
|  | significant | not-significant |  |

**Table 2. Overlap of autosome-wide SNPs between HWE meta-analyses of 10 groups.**
|  |  |  |  |  |
| --- | --- | --- | --- | --- |
| **HWE** | significant | 14,982 (140) | 7 (3) | 14,989 (143) |
| **Heterogeneity**<br>**(reliable)** | not-significant | 15,956 (115) | 3,346,256<br>(1,347,737) | 3,362,212<br>(1,347,852) |
| **Total** |  | 30,938 (255) | 3,346,263<br>(1,347,740) | 3,377,201<br>(1,347,995) |

Similarly, Figure 3 shows the distribution of HWE p-values on the X chromosome. Figure 4a shows the HWE distribution from the omnibus meta-analysis of SNPs with MAF>=5% in all groups. Across all 10 groups, there were 5,381 PAR1 and 277 PAR2 SNPs, as well as 97,225 NPR SNPs in the 5 female groups. Figure 4b shows the corresponding HWE heterogeneity results. In contrast to the autosomes, the PARs exhibited greater similarity in HWD patterns across super-populations than between sexes, essentially because males showed higher HWD near the PAR–NPR boundaries.

**Figure 3.**
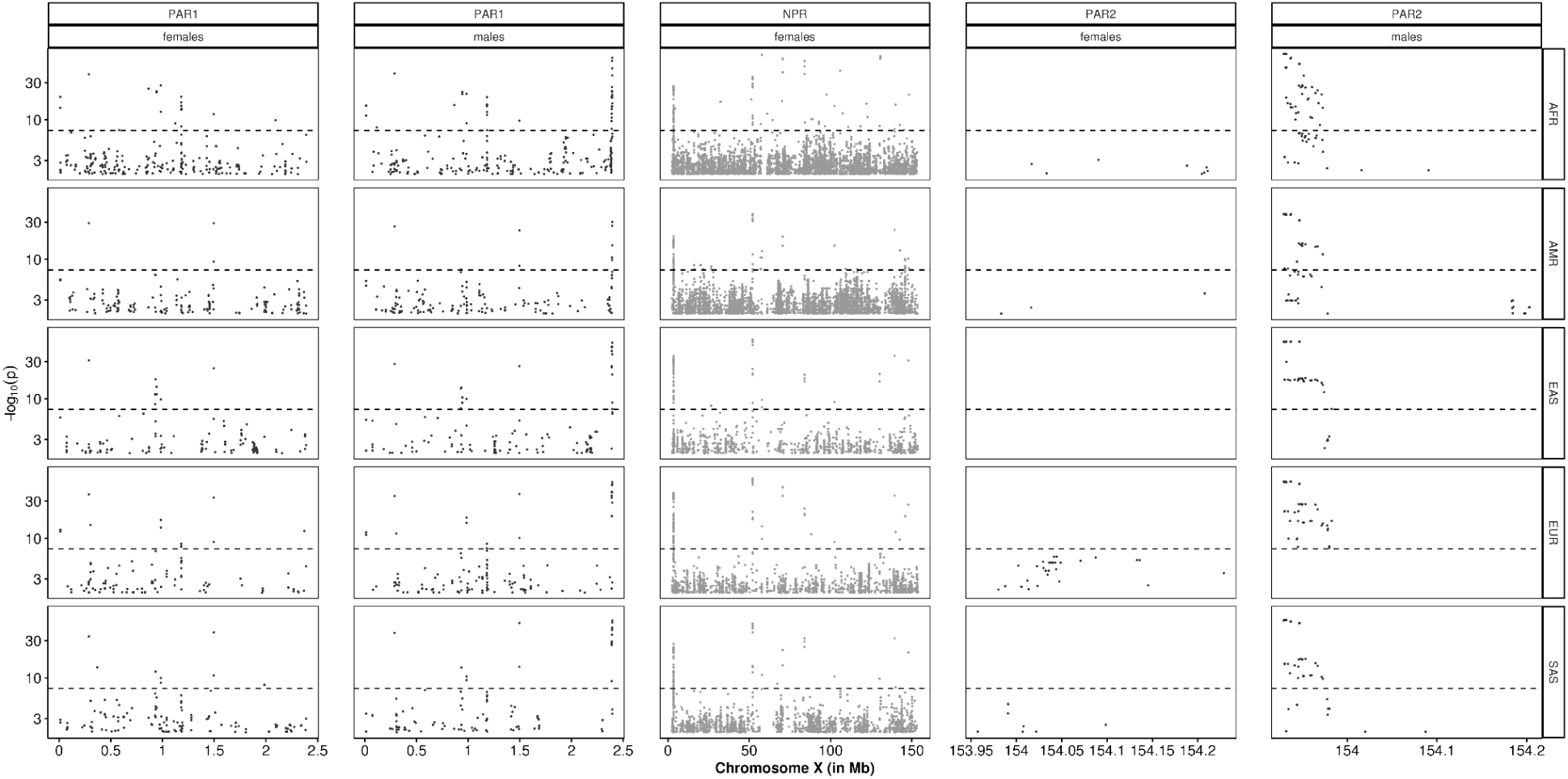
Manhattan plots for X-chromosome HWE in each of the 10 groups. HWE results from five super-populations and two sexes are shown for chrX SNPs with MAF>=5%. NPR data is only for females. Dashed line represents the genome-wide significant threshold of p=5e-8. Y-axis is presented on a log10 scale for SNPs with HWE p-value<0.01.

**Figure 4.**
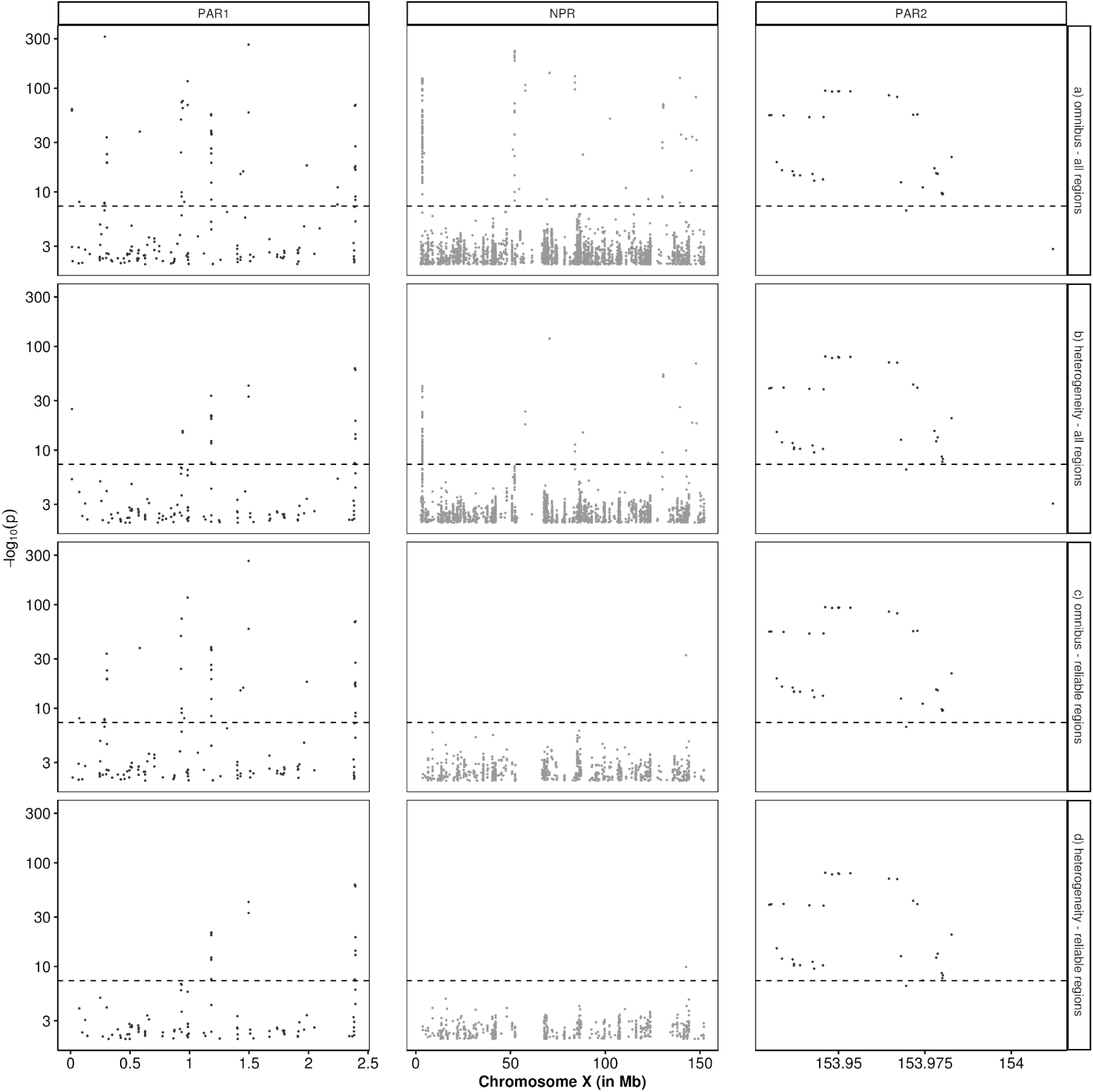
Manhattan plots for X-chromosome HWE meta-analysis of 10 groups. Meta-analysis results from five super-populations and two sexes are shown for chrX SNPs which have MAF>=5% in all groups: a) omnibus and b) HWE heterogeneity results for all regions (PAR1=5,381, PAR2=277, NPR=97,225), c) omnibus and d) HWE heterogeneity results for reliable regions (PAR1=4,473, PAR2=252, NPR=26,890). NPR data is only for females. Genomic features were not available for PARs except difficult-to-sequence regions. Dashed line represents the genome-wide significant threshold of p=5e-8. Y-axis is presented on a log10 scale for SNPs with meta-analysis p-value<0.01.

In the omnibus meta-analysis, 1% SNPs on the autosomes were in HWD, compared to 1.5% and 0.3% SNPs on the PARs and NPR, respectively. In the HWE heterogeneity meta-analysis, 0.44% SNPs on the autosomes were in HWD, compared to 0.97% and 0.07% SNPs on the PARs and NPR, respectively. Accompanying Figures 1-4, Figures S1-S4 show the corresponding QQ plots and histograms of HWE testing p-values.

Additionally, a heterogeneity test measuring sex differences in HWD (sdHWD) within each super-population yielded 80 unique genome-wide significant SNPs (24 in autosomes, 20 in PAR1 and 36 in PAR2); all 80 SNPs had MAF>=5% in both sexes of a super-population.

### Effect of genomic features on HWD

Certain regions of the genome are prone to misalignment in short-read sequencing, leading to genotyping errors. We evaluated the effect of these genomic features with predefined positions on HWD patterns and created a list of “reliable regions” by restricting the genome to positive features and removing the regions with negative features (see Methods).

Figure 5 shows the enrichment of negative genomic features in autosomal SNPs with HWD. Over half of the SNPs in three negative features (difficult-to-sequence regions, segmental duplications and centromere/satellite regions) had significant HWD, even though they constituted less than 10% of the autosomal SNPs. In contrast, less than a third of SNPs in the positive feature (short-read accessibility mask) had significant HWD, even though they covered 96% of the autosomes. When the negative features were combined, they accounted for 96% of SNPs with significant HWD. Table S2 describes the number of SNPs and the 𝝌²_1_ test for each of these features.

**Figure 5.**
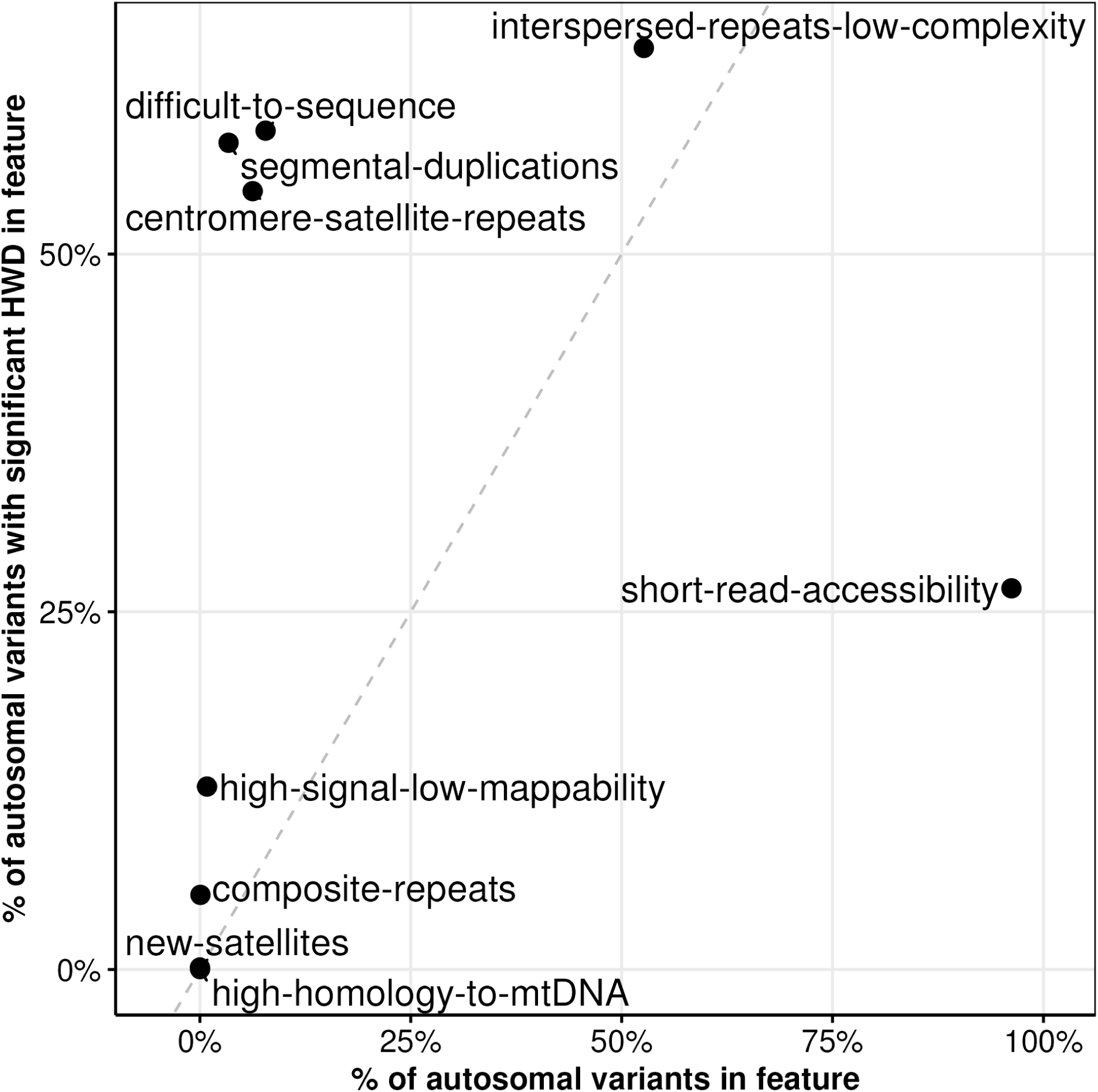
HWD in autosome-wide genomic features. Scatter plot of genomic features compares the autosomal percentage of SNPs in that feature (out of 3,377,201 SNPs) to the percentage of SNPs with significant HWD in that feature (out of 30,938 SNPs). Two genomic features (“assembly-gaps” and “telomere”) had no overlaps with the analyzed genome and hence are not displayed here. Dashed line represents the identity line.

Figure 2c shows the distribution of HWE p-values from the omnibus meta-analysis after restricting positions to reliable regions in the autosomes (1,347,995 SNPs, which is 40% of unrestricted autosomal SNPs analyzed above). Restricting positions reduced the autosomal SNPs with genome-wide significant HWD from 1% to 0.02% (from 30,938 to 255 SNPs). Figure 2d shows the corresponding HWE heterogeneity results.

There were 56 significant SNPs with no adjacent SNPs within a 100kb-interval that were also genome-wide significant; however, a few clusters of significant SNPs emerged after the omnibus meta-analysis.

In parallel, Figure 4c shows the distribution of HWE p-values from the omnibus meta-analysis after restricting positions to reliable regions on the X chromosome (26,890 in NPR, 4,473 in PAR1 and 252 in PAR2). Restricting PAR positions increased the proportion of SNPs in HWD from 1.54% to 1.59% because the only available genomic feature for PARs was difficult-to-sequence regions (the other features were not available for the PARs). However, restricting NPR positions to reliable regions reduced the SNPs in HWD from 0.3% to a single SNP (rs859902). Figure 4d shows the corresponding HWE heterogeneity results.

### Autosomal SNPs with HWD in reliable regions

The 255 autosomal SNPs with HWD in reliable regions (as shown in Figure 2c) fell into 90 distinct 100 kb non-overlapping intervals. Table S3 provides genomic details for the 256 SNPs (255 on autosomes and 1 on NPR) after lifting over to GRCh38 for functional annotations. Five autosomal SNPs were not present on GRCh38, and an additional 15 had failed QC in the original work, which was aligned to GRCh38 (2). Table S4 provides the NHGRI-EBI GWAS catalog results for the 10 SNPs with GWAS p<5e-8 (see Table S5 for a summary of the number of studies associated with each SNP). Three out of these 10 SNPs were in the major histocompatibility complex (MHC) region on 6p21 and were associated with multiple blood cell traits. All 10 SNPs were found in studies primarily based on European populations, likely reflecting the gross over-representation of European populations in the GWAS catalog (25).

Table S6 provides group-wise allele counts and HWD results for the 256 SNPs, and Table S7 reports their meta-analysis results. Of the 255 autosomal SNPs, 140 exhibited significant heterogeneity in HWD across the 10 groups (Table 2). Figure 6 shows the HWD delta across groups, generally showing consistency, but with exceptions, such as at the chr14 region described below. Of the 256 SNPs, 200 have a negative HWD delta in all groups (Figure S5), indicating an excess of heterozygotes. 53 of 256 SNPs had significant HWD within all groups, and another 35 SNPs had significant HWD only in the AFR male and female groups (Figure S6). No autosomal sdHWD SNPs were observed in reliable regions. After removing difficult-to-sequence regions, no PAR1 SNPs and only one PAR2 SNP remained; this SNP was found in the EUR super-population and showed a significant sdMAF (Table S8).

**Figure 6.**
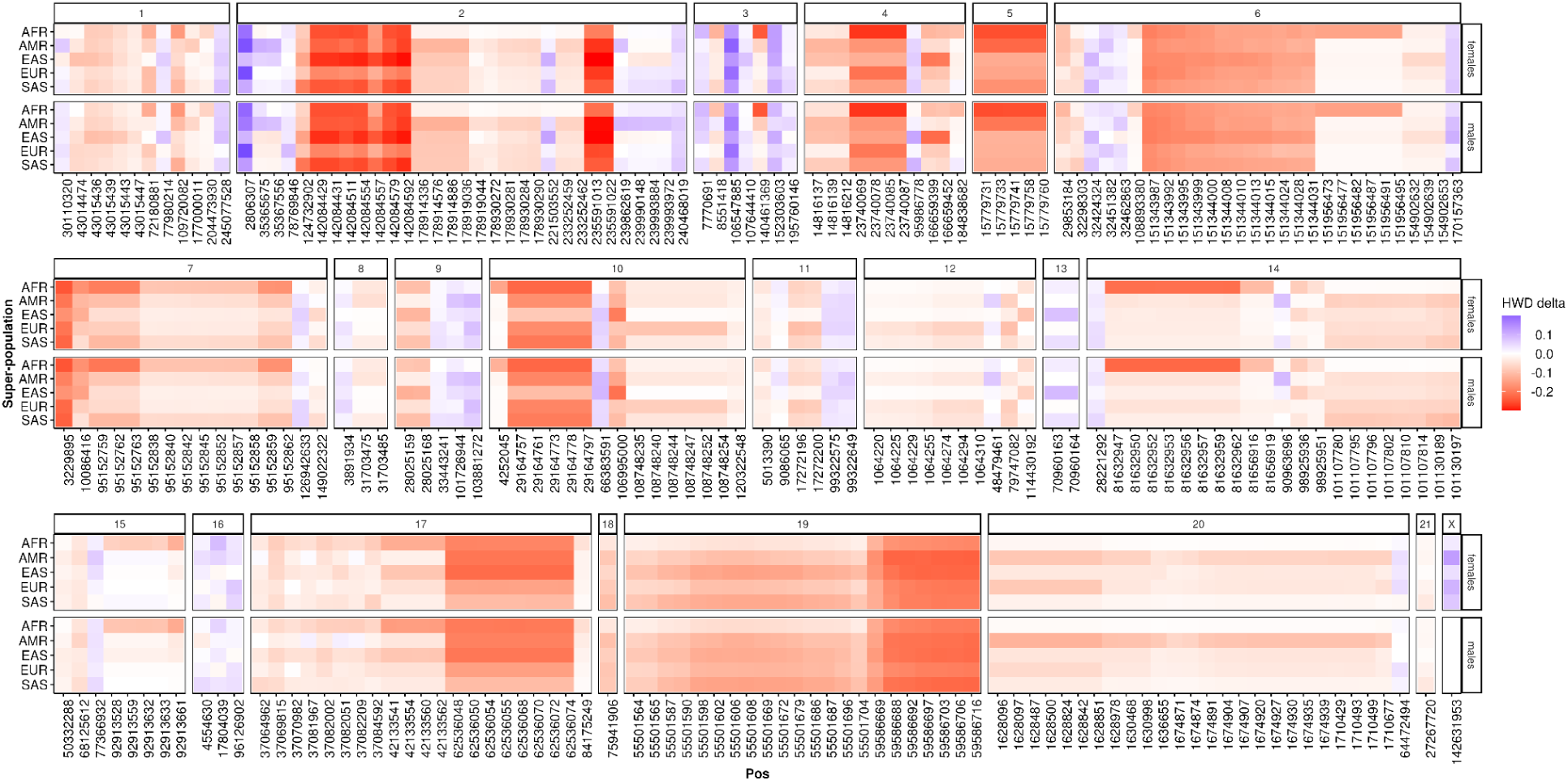
Heatmap of HWD delta for 256 SNPs with significant HWD. HWD delta is shown by a signed color scale for each of the 10 groups at the 256 SNPs (Table S7). Panels represent chromosomes and genomic location with each chromosome is arranged in ascending order on the X-axis. Red gradient represents negative delta, which means excess of heterozygotes. See companion Figure S5 for non-gradient colors.

Notably, an interesting sequence of eight SNPs was observed over a 15-bp region (chr14:81,632,947-81,632,962 at 14q31.3), where the reference and alternate sequences were reverse complements of each other (Figure S7). HWD at this region was driven by heterozygosity excess in both sexes of the AFR super-population, where over 90% of individuals were heterozygous at all 8 SNPs (Figure 7). In contrast, non-AFR populations showed no significant HWD at this region (Table S7). These SNPs passed QC in the original work, which was aligned to GRCh38 (2). Allele bias at this region for the AFR super-population is high both in v2 (Figure S8) as well as in GRCh38 (Figure S9) with estimates of 5:1 allele depth across sexes and SNPs. This could indicate unrecognized repetitive variation at this region in AFR samples, such as a VNTR making genotyping challenging. Additional quality parameters in v2 are provided in Figure S10.

**Figure 7.**
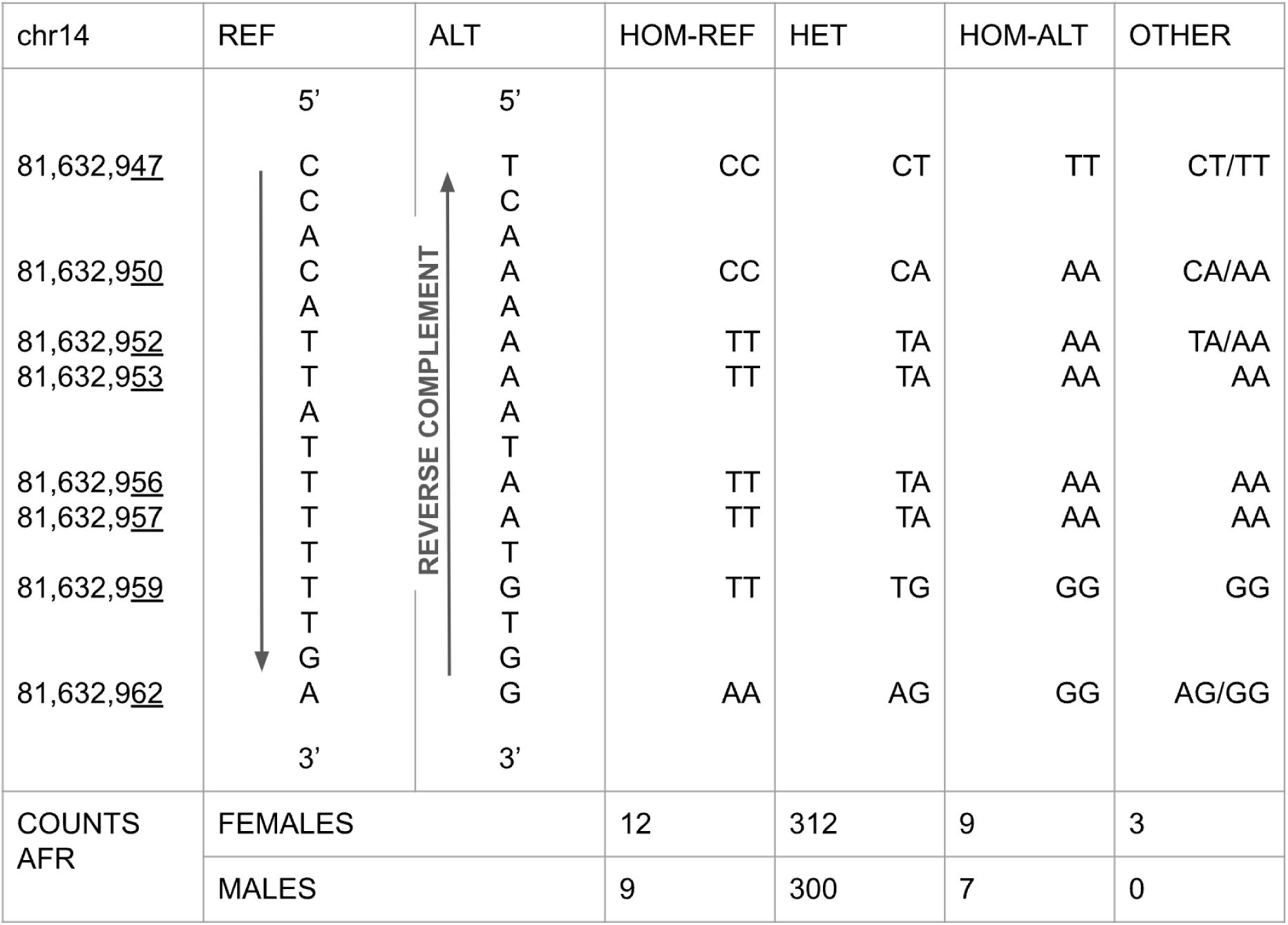
AFR genotype counts at the chr14 locus. The alternate sequence over a 15bp region at chr14:81,632,947-81,632,962 is the reverse complement of its reference sequence (see Figure S5). This region consists of 8 SNPs where HWD was observed (see Table S7). These 8 SNPs have high LD and excess heterozygotes in both male and female AFR samples, as shown by the haplotypes formed by their diploid genotype counts.

### Multivariate genomic feature model for autosomal HWD identified by the omnibus test prior to restricting to the reliable regions

Many features associated with HWD (Table S2) showed inter-correlation and overlaps (Figure S11-12). As expected, “short read accessibility”, a positive feature, was negatively correlated with others. Additionally, features with fewer SNPs, such as “new satellites” or “high homology to mtDNA”, showed little correlation with others.

Five features were selected based on their importance (Figure S13) and AIC to be used in a multivariate model. To account for linkage disequilibrium among nearby SNPs, we retained the first SNP in each 100 kb non-overlapping interval for both the training set (odd autosomes: 13,524 SNPs of which 190 had significant HWD) and the test set (even autosomes: 13,358 SNPs of which 103 had significant HWD). The model predicted HWD significance status with high accuracy (AUC is 99.4%; Table S9 and Figure S14).

### Comparison of autosomal and NPR HWD in recently-admixed populations

Because recent admixture can result in HWD, we examined this. In recently-admixed populations (ASW and ACB) from the AFR super-population, we examined HWE separately for autosomes (79 females and 71 males) and the chrX NPR (79 females) stratified by MAF (SNPs in reliable regions at each MAF bin: 339,046 in 0.1<=MAF<0.2, 330,291 in 0.2<=MAF<0.3, 315,586 in 0.3<=MAF<0.4 and 308,859 in MAF>=0.4). Inspection of QQ plots (Figure S15) revealed that SNPs in NPR did not deviate from HWE, whereas SNPs in most MAF bins in the autosomes had inflated HWD (except 0.1<=MAF<0.2). Inspection of HWD delta statistics (Figure S16) in the significant SNPs (p<5e-8) reveals that most SNPs exhibit a heterozygote excess (Table S10).

## Methods

### Data

We used high-coverage whole genome sequences from the 1000 Genomes Project aligned to T2T-CHM13v2.0 using XYalign (5), which we refer to as “v2”. To our knowledge, this data has not been subjected to quality control based on HWE thresholds. SNPs were filtered after applying VQSR (v4.1.9.0) using known sites lifted over from GRCh38 (as described at https://github.com/schatzlab/t2t-chm13-chry). We performed further quality control on the data by removing up to second-degree relatives and selecting bi-allelic SNPs with non-missing genotypes. We analyzed both autosomes (chromosomes 1-22) and sex chromosomes (chromosomes X and Y). The two pseudo-autosomal regions (PAR1 and PAR2) of sex chromosomes were studied in all individuals. The non-pseudo-autosomal region (NPR) of the X chromosome was only studied in females, as the concept of HWE does not apply to males in NPR.

For the primary HWD analyses, we used 2,490 unrelated samples (1,263 females and 1,227 males) from v2 for our study. We divided these individuals by super-population and sex to form 10 groups (5 super-populations x 2 sexes) for all analyses (except the recent-admixture analysis), which was conducted on SNPs with MAF>=5% in all groups.

For the secondary analysis of HWD in recently-admixed populations, we used 150 unrelated samples (79 females and 71 males) from v2 that belonged to the ASW (African Ancestry in Southwest US) and ACB (African Caribbean in Barbados) populations. Here, due to a smaller sample size, we analyzed SNPs with MAF>=10% in the 150 samples.

### Quantifying HWD

We quantify departure from Hardy–Weinberg equilibrium (HWE), i.e., Hardy-Weinberg disequilibrium (HWD), as δ, the frequency difference between one of the observed homozygous genotypes and its expectation under HWE in a given population/group (26). Specifically,

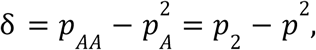

where 𝑝*_AA_* is the 𝐴𝐴 genotype frequency and 𝑝*_A_* the 𝐴 allele frequency; δ = 0 indicates HWE and δ < 0 indicates excess heterozygosity.

### Test statistics for one group

The test statistic takes the general form,

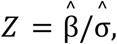

where β and σ are context-dependent parameters, and 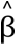 and 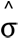 are their estimates.

When testing for HWD,

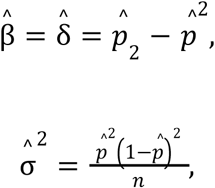

and the resulting test statistic 𝑇_𝐷_ is,

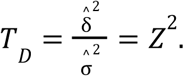

Under the null hypothesis of HWE, 𝑇*_D_* is asymptotically chi-squared distributed with one degree of freedom, 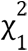. Although 𝑇*_D_* does not have the usual Pearson chi-square form, it is equivalent to the traditional test (27). Males were excluded from NPR analyses because HWE does not apply to NPR variants in males.

### Meta-analyzing HWD across groups

For group 𝑘, 𝑘 = 1, … 𝐾, let

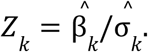

To test for HWD across the 𝐾 groups, the classical meta-analysis is suboptimal because it uses the weighted average of 𝑍*_k_* and is therefore sensitive to the direction of β’s (i.e. of δ’s). A priori, one group may show excess heterozygosity (δ < 0) while another shows depletion (δ > 0). We therefore use an omnibus meta-analysis aggregating 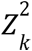 (28,29). The resulting statistic 𝑇*_M_* appears below and is asymptotically 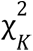 distributed under the null of HWE in all 𝐾 populations.

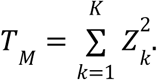

### Heterogeneity tests

To assess between-group differences in effect size (β), we use Cochran’s 𝑄,

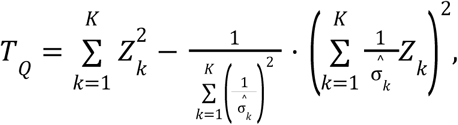

which is asymptotically 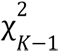 distributed under the null of no heterogeneity. We apply 𝑇*_Q_* to evaluate HWD heterogeneity across the 10 groups (five superpopulations and two sexes).

For each super-population, we first test for sex differences in HWD (sdHWD). We then compare MAFs between the sexes (sdMAF; (17)), noting that the magnitude of HWD δ depends on MAF: − 𝑝^2^ ≤ δ ≤ 𝑝(1 − 𝑝) (27). We note that when 𝐾 = 2 as in sdHWD and sdMAF analyses, 𝑇*_Q_* reduces to the familiar two-sample comparison with group-specific variance estimates,

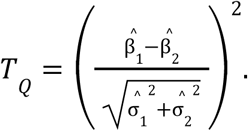

For sdMAF on autosomal and PAR SNPs we use,

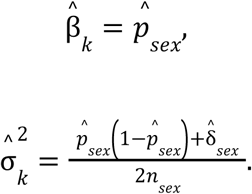

The sdMAF statistic for NPR SNPs differs slightly (17); we omit it here because we analyze only SNPs for which HWD testing is possible in males.

### Genomic features

Short-read next-generation sequencing has difficulties in correctly calling certain regions of the genome, which can impact HWE results. Therefore, we created a list of “reliable regions” using predefined annotations. We used several published annotations available for the T2T-CHM13 reference genome provided by a) the “Telomere-to-Telomere” (T2T) Consortium: i) centromere/satellite repeats (30), ii) composite repeats (31), iii) segmental duplications (32), iv) new satellites (30), v) interspersed repeats and low complexity regions (31); b) Ogata et al., 2023: vi) high signal and low mappability regions, vii) regions of high homology to mtDNA (nuclear mitochondrial DNA), viii) assembly gaps, ix) telomeres; c) Genome-In-A-Bottle: x) difficult to sequence regions [Citation error]; d) Illumina: xi) short-read accessibility mask (19). See Table S11 for more details.

All features except for the short-read accessibility mask are “negative” features. Thus, we created a set of “reliable regions” by removing them from the “positive” short-read-accessibility mask. Two genomic features (“assembly gaps” and “telomeres”) were a subset of other features and did not have unique SNPs. No genomic features were available for PARs, except “difficult to sequence regions”.

### Variant annotations

We used the Entrez Programming Utilities (edirect v22.0) to query dbSNP (build 156) to find rsIDs, variant class, and clinVar significance of a SNP.

We used the Ensembl Variant Effect Predictor REST API (v15.8; https://rest.ensembl.org/vep/human/id/{rsid}) to find the most severe consequence of a SNP.

We queried the GWAS catalog APIs with different parameters and merged all results in order to be comprehensive: i) https://www.ebi.ac.uk/gwas/rest/api/singleNucleotidePolymorphisms/{rsid}/associations?projection=associationBySnp, ii) http://www.ebi.ac.uk/gwas/summary-statistics/api/associations/{rsid}?p_upper=5e-8, iii) http://www.ebi.ac.uk/gwas/summary-statistics/api/associations/{rsid}?size=100. GWAS catalog was queried in June 2024 at which time it was mapped to GRCh38.p14 and dbSNP build 156.

### Predictive model

We created a logistic regression model using five binary genomic features to predict a binary HWD significance status on the autosomes. We selected the features using the MASS::stepAIC (v7.3-59) function in R (v4.3.0). SNPs in linkage-disequilibrium (LD) can unduly affect the model, which are often pruned in an ancestry-specific manner.

However, given our multi-ancestry data, we opted for a simpler approach where we selected the first SNP in each 100 kb window on every autosome since LD typically decays at that physical distance (1). We assigned all odd autosomes to the training set and all even autosomes to the test set.

## Discussion

The common strategy of relying on statistical significance alone in HWE tests ignores the direction and magnitude of the deviation (33) as well as the sample size. These are important when the data has multiple groups. Some methods, such as RUTH (Robust Unified Test for Hardy–Weinberg Equilibrium) (34) and others (28), account for heterogeneous groups. However, there is still a need for methods that not only consider the context of HWE in a multi-group setting but also quantify the degree of deviation to allow for the comparison and testing of differences in HWD across groups. Combining and contrasting HWD across groups is a complex task, particularly when the direction of effect (δ) varies between groups.

We chose the 1KGP because it is a relatively large, publicly available and widely used resource. The vast majority of analyses of 1KGP, including previous HWE analyses (9), utilized the Phase 3 data, which relies primarily on low-coverage whole-genome sequencing. Here, instead, we use high-coverage whole-genome sequence data that has been aligned to T2T-CHM13v2.0, which has previously been shown to improve variant calling (4). We performed sex- and super-population-stratified HWD analysis on this data. We included sex stratification in our analysis because recent work (12,17,21,35) has shown allele frequency differences between sexes at a small proportion of SNPs. When we examined sex differences in HWD, we identified only 80 SNPs with significant sdHWD, the majority of which were located in PARs, and only one survived the restriction to reliable regions. Understanding the varied contributors to HWD across populations is especially noteworthy, as neglecting such influences can result in the inappropriate application of quality control strategies (13,35,36).

Since HWD delta (δ) is restricted by MAF, we applied a strict MAF threshold before analyzing our data, which may not be necessary in larger studies. For example, some recent works examining heterozygosity excess (37) or homozygosity deficit (38) have focused predominantly on autosomal SNPs with low MAF in GRCh38-aligned data sourced from multiple studies and technologies. Abramovs et al., 2020 restricted their analysis to protein-coding SNPs in gnomAD 2.1.1 with MAF<0.05. Oddsson et al., 2023 exclusively analyzed European populations from multiple cohorts and included both imputed and sequenced data to specifically study protein-altering SNPs. Another important limitation of our work is that we only examined bi-allelic SNPs with no missing data that did not overlap with indels. Although multi-allelic extensions of HWE have been developed (e.g. (39)), for example, for microsatellites, we did not implement them here for simplicity.

In our work, we emphasize the effect of unreliable regions on HWD. We found that, in the autosomes, restricting positions to reliable regions reduced the SNPs in HWD from 1% to 0.02%. In fact, three negative genomic features (difficult-to-sequence regions, segmental duplications, and centromere/satellite regions) that encompassed less than 10% of all autosomal SNPs accounted for a large proportion of significant HWD results. This is also supported by the multivariate model results, where these three features are the most significant predictors of HWD. No genomic features were available for PARs, except “difficult to sequence regions”.

HWD patterns in all groups are similar, with deviation typically occurring at telomeres and around centromeres in autosomes (Figure 1; note that chromosomes 13-15, 21, 22 are acrocentric). One very notable region with population-specific deviation was at chr14q31.3, where HWD was observed in the AFR super-population in 8 SNPs over a 15bp-region. Interestingly, the reference and alternate haplotypes are reverse complements of each other (Figure S7). This raises the possibility that reads from this region are not being called properly, which can be due to several reasons. Sequence misalignment can result in erroneous genotypes. Technical artifacts like sample contamination can lead to systematic genotyping errors (40,41); Byrska-Bishop et. al., 2022 used a threshold of 2% to eliminate contaminated samples. Long-read sequencing technology can also cause technical artifacts such as foldbacks (duplicated inversions) (42).

However, the high linkage disequilibrium between these 8 SNPs is not limited to the data we have analyzed but has also been observed in the pangenome, which included long and linked-read sequencing data for some samples from 1KGP (see Resources and Figure S17; (43). Since there were only 24 individuals of African ancestry in the pangenome data, we did not perform statistical analysis of HWD. Moreover, the reference sequence is identical in T2T CHM13v2.0 and GRCh38, even though T2T CHM13v2.0 is of European origin (3) whereas GRCh38 is predominantly of African-European origin (44,45). Also, the ratio of allele depths for these 8 SNPs follows the same pattern in data aligned to both T2T CHM13v2.0 and GRCh38 (Figure S9). This region has no alternate contigs in GRCh38.

We note that this region exhibits a high density of variation in dbSNP (build 157, Figure S18) and overlaps with two multi-nucleotide variants (MNVs, rs796238959 and rs770667319, on GRCh38.14; Figure S19). rs796238959 (chr14:87412183-87412205; ss1808029058) is derived from the Western African Pygmy population (46). rs770667319 (chr14:87412188-87412200; ss1576679280) is derived from the GenomeDenmark project (47). MNVs can give rise to alignment challenges (48), and impact variant calling. These SNPs are intronic in all the four reported transcripts of LINC02296 (long intergenic non-protein coding RNA gene). We also note that the unusual structure of reference and alternate haplotypes at this region resembles an inversion heterozygote (49). Moreover, given that the allele depth ratio in AFR super-population is 4.8, it is possible that this region contains structural variation, but it has not been reported (Table S11).

Separately, we examined the impact of recent-admixture on HWD in the chrX. Theoretically, it takes more generations for HWE to be achieved on chrX compared to the autosomes (23,24), but this has not been tested. Therefore, we used two recently-admixed populations (African Ancestry in Southwest US [ASW], African Caribbean in Barbados [ACB]) in our analysis. We compared HWD in NPR and autosomes stratified by MAF. Contrary to the theoretical expectation, our results indicate that whereas 80 of 1,347,995 autosomal SNPs deviate from HWE, none of the 26,890 NPR SNPs do. As expected, we observed an excess of heterozygotes in the SNPs that deviate from HWE (Figure S16). In addition, African American populations have been shown to have higher African ancestry on chrX owing to the sex bias in ancestry contributions brought about by asymmetric mating (50). Our analysis has limited scope due to small sample size, especially so on the chrX, and deserves attention in future larger populations including other admixed groups.

A general limitation of this work is that the sample size is modest and SNPs were restricted (biallelic SNPs with MAF>=5% and no missingness). HWD patterns for other variant types, which were not examined here, may differ.

## Resources

### WGS data (‘v2’)

https://s3-us-west-2.amazonaws.com/human-pangenomics/index.html?prefix=T2T/CHM13/assemblies/variants/1000_Genomes_Project/chm13v2.0/unrelated_samples_2504/

### WGS data (GRCh38)

http://ftp.1000genomes.ebi.ac.uk/vol1/ftp/data_collections/1000G_2504_high_coverage/working/20201028_3202_raw_GT_with_annot/

### 1KGP ancestries

https://projecteuclid.org/journals/annals-of-applied-statistics/volume-17/issue-2/Leveraging-HardyWeinberg-disequilibrium-for-association-testing-in-case-control-studies/10.1214/22-AOAS1695.short

### UCSC genome browser result for chr14 locus on GRCh38 (Multiple alignments of ‘Human Pangenome - HPRC’ track can be turned on)

https://genome.ucsc.edu/cgi-bin/hgTracks?db=hg38&lastVirtModeType=default&lastVirtModeExtraState=&virtModeType=default&virtMode=0&nonVirtPosition=&position=chr14%3A87412182%2D87412206&hgsid=2500872119_ZhA4jGydieyrY2C5Mn02OWsx0S0p

### UCSC genome browser result for chr14 locus on T2T CHM13v2.0

https://genome.ucsc.edu/cgi-bin/hgTracks?db=hub_3671779_hs1&lastVirtModeType=default&lastVirtModeExtraState=&virtModeType=default&virtMode=0&nonVirtPosition=&position=chr14%3A81632947%2D81632962&hgsid=3326470641_djDKIWfSW1n98DvnDjd76A7gmlSu

## Acknowledgements

We acknowledge the “Telomere-to-Telomere” (T2T) Consortium for the creation of some of the datasets used in this manuscript.

