## Supplementary Figures for "Assessing Hardy-Weinberg Equilibrium in T2T-aligned 1000 Genomes Project"

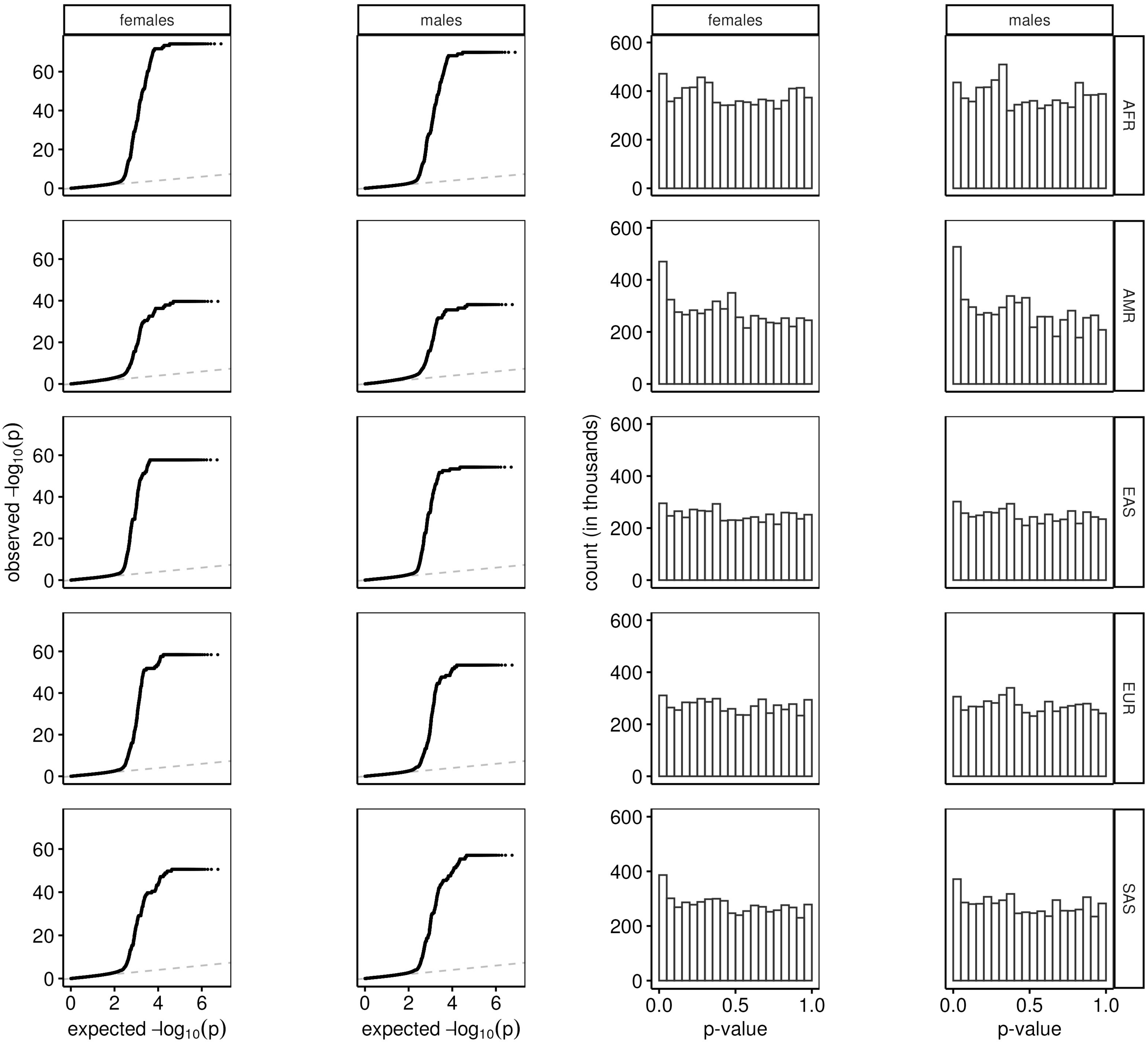

females
males
females
males
AFR
AMR
EAS
EUR
SAS
observed -log10(p)
expected -log10(p)
count (in thousands)
p-value
0
20
40
60
0
2
4
6
0
200
400
600
0.0
0.5
1.0

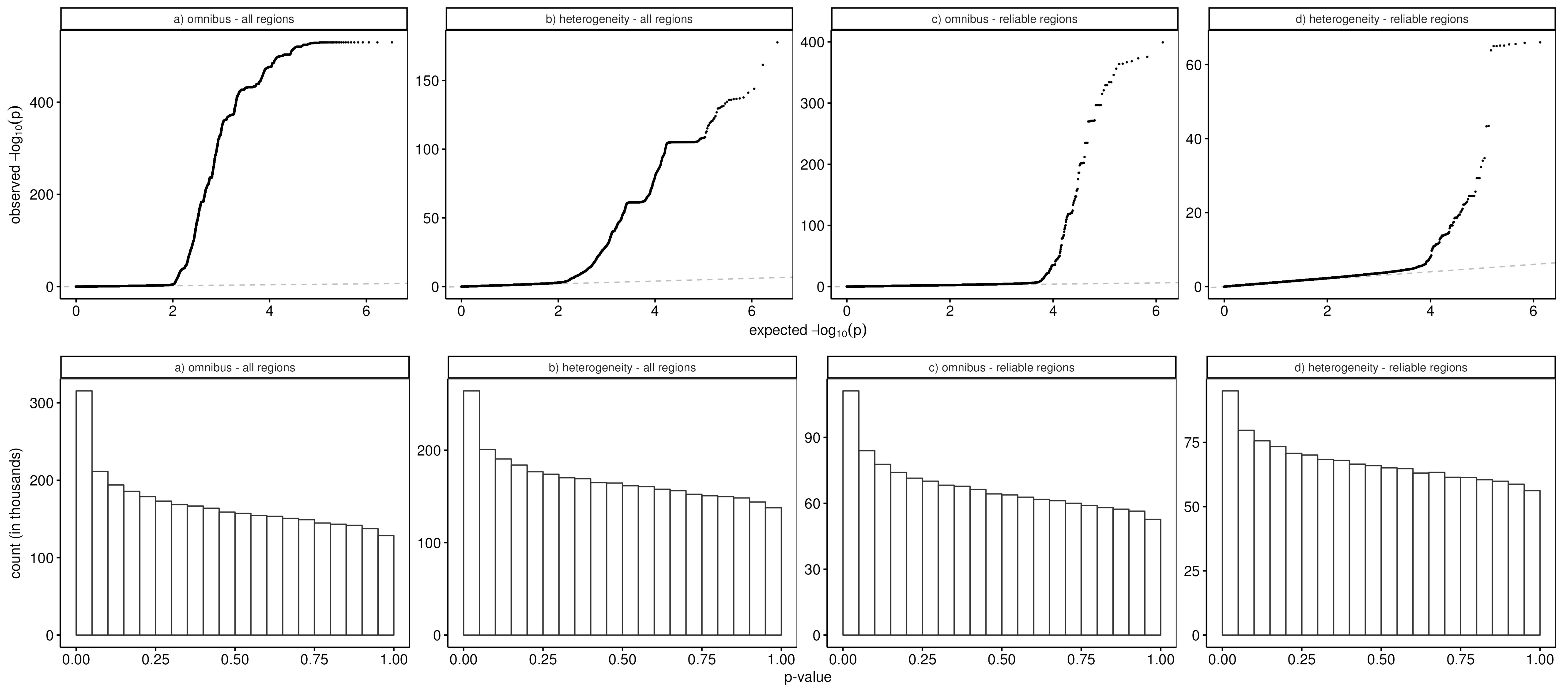
a) omnibus - all regions
b) heterogeneity - all regions
c) omnibus - reliable regions
d) heterogeneity - reliable regions
observed $-\log_{10}(p)$
expected $-\log_{10}(p)$
a) omnibus - all regions
b) heterogeneity - all regions
c) omnibus - reliable regions
d) heterogeneity - reliable regions
count (in thousands)
p-value

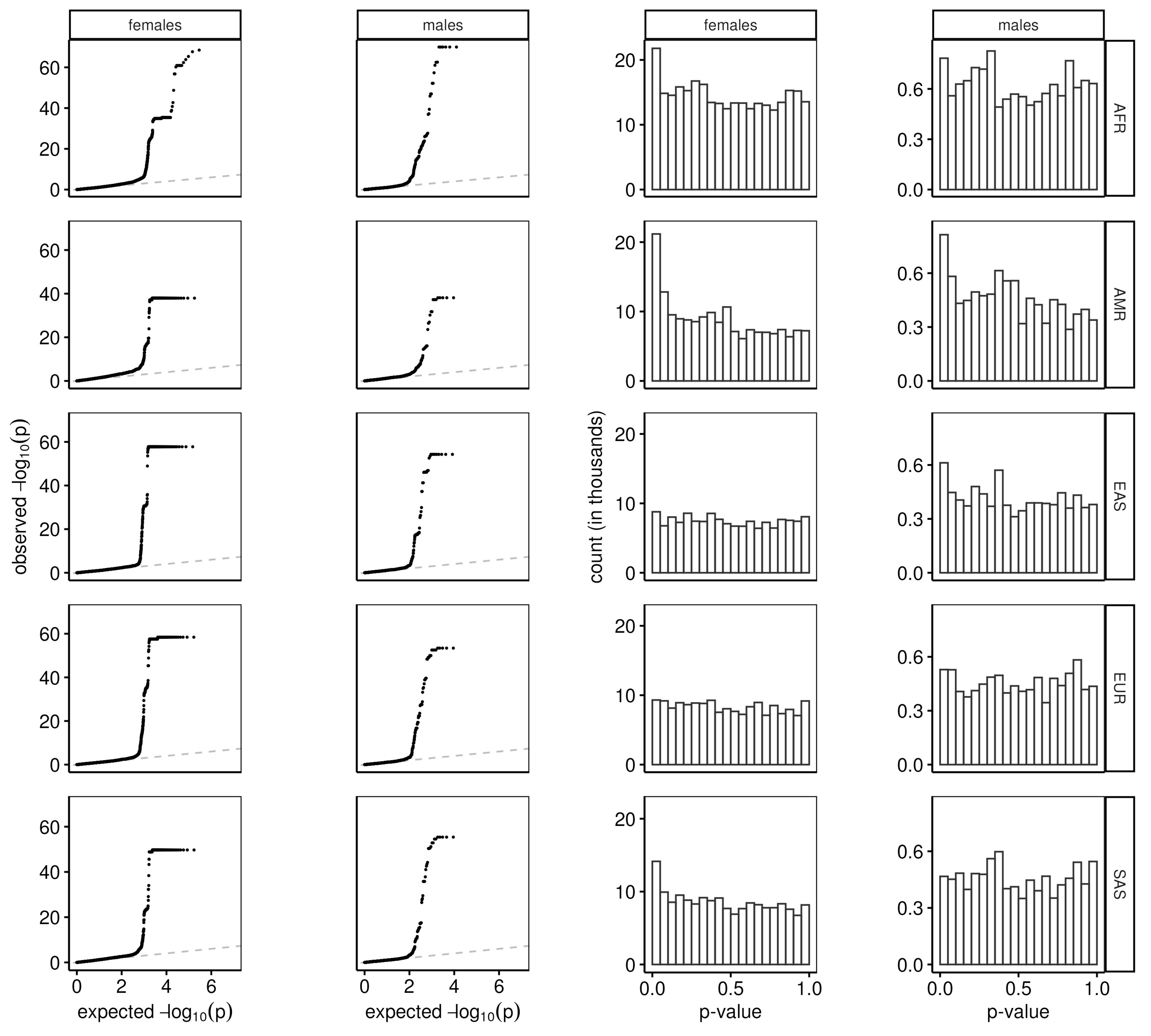
females
males
females
males
AFR
AMR
EAS
EUR
SAS
observed $-\log_{10}(p)$
expected $-\log_{10}(p)$
count (in thousands)
p-value

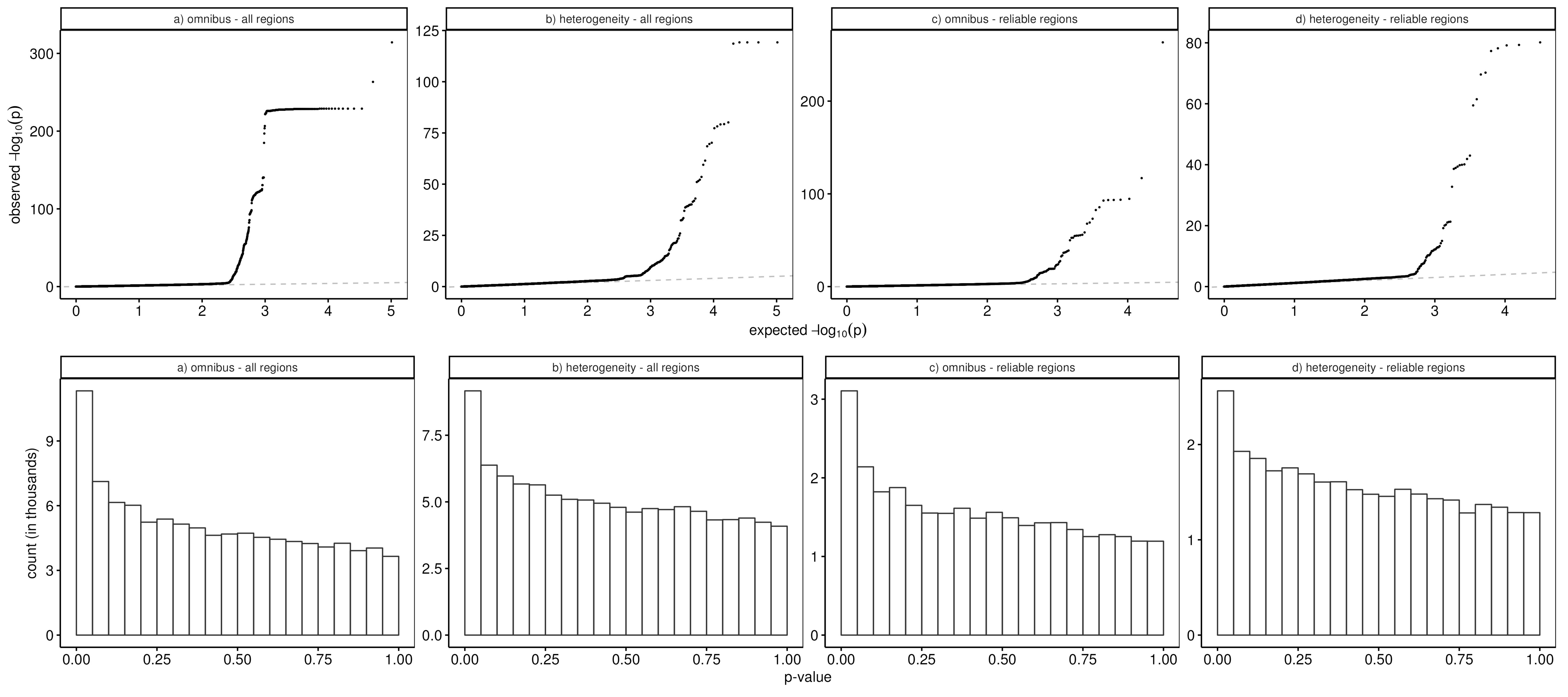
a) omnibus - all regions
b) heterogeneity - all regions
c) omnibus - reliable regions
d) heterogeneity - reliable regions
observed −log10(p)
expected −log10(p)
count (in thousands)
p-value

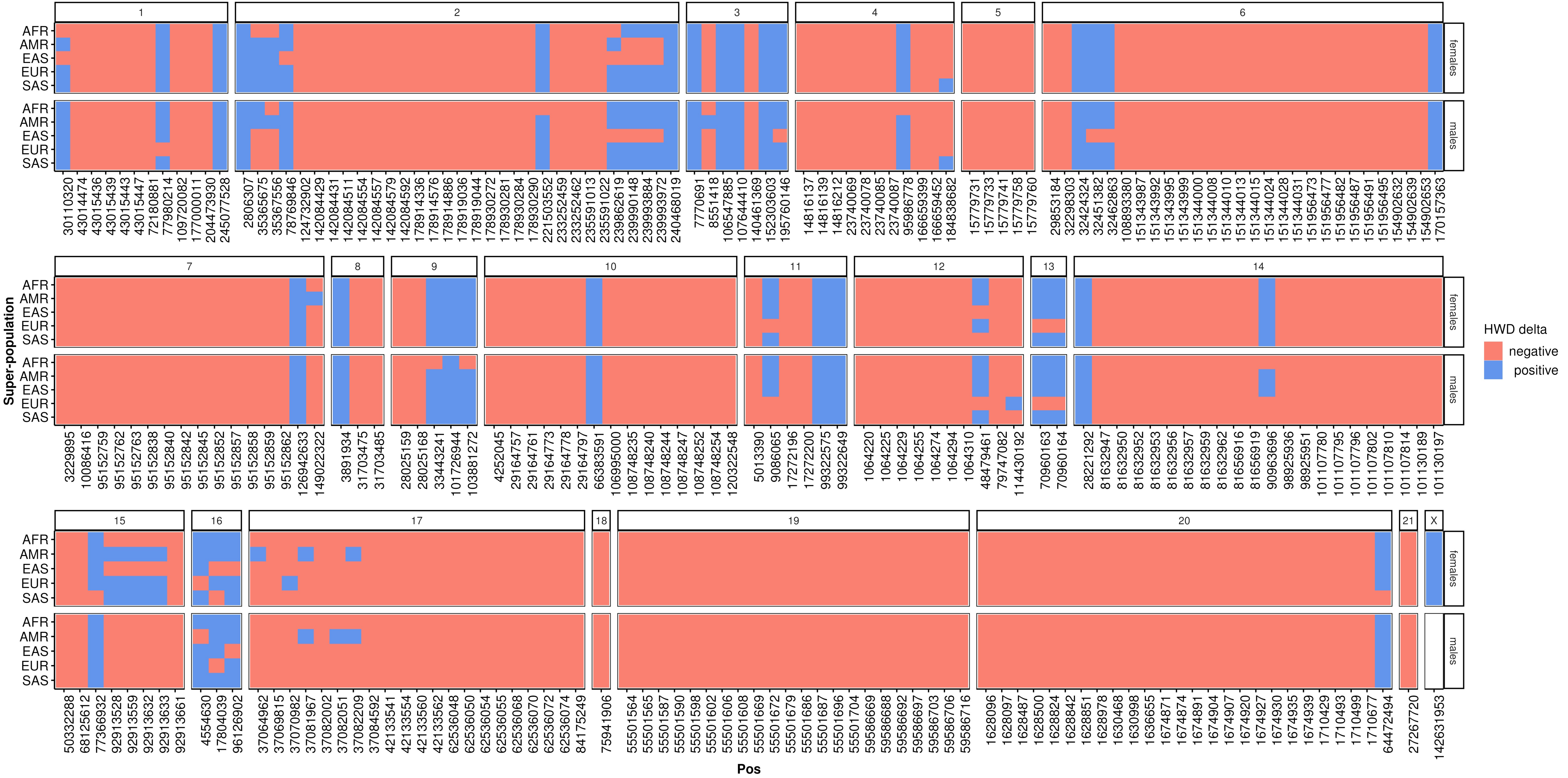

X
AFR
AMR
EAS
EUR
SAS
females
males
Super-population
Pos
HWD delta
negative
positive
30110320 43014474 43015436 43015439 43015443 43015447 72180881 77980214 109720082 177000011 204473930 245077528
2806307 35365675 35367556 78769846 124732902 142084429 142084431 142084511 142084554 142084557 142084579 142084592 178914336 178914576 178914886 178919036 178919044 178930272 178930281 178930284 178930290 221503552 233252459 233252462 235591013 235591022 239862619 239990148 239993884 239993972 240468019
7770691 8551418 106547885 107644410 140461369 152303603 195760146
14816137 14816139 14816212 23740069 23740078 23740085 23740087 95986778 166659399 166659452 184838682
15779731 15779733 15779741 15779758 15779760
29853184 32298303 32424324 32451382 32462863 108893380 151343987 151343992 151343995 151343999 151344000 151344008 151344010 151344013 151344015 151344024 151344028 151344031 151956473 151956477 151956482 151956487 151956491 151956495 154902632 154902639 154902653 170157363
3229895 10086416 95152759 95152762 95152763 95152838 95152840 95152842 95152845 95152852 95152857 95152858 95152859 95152862 126942633 149022322
3891934 31703475 31703485
28025159 28025168 33443241 101726944 103881272
4252045 29164757 29164761 29164773 29164778 29164797 66383591 106995000 108748235 108748240 108748244 108748247 108748252 108748254 120322548
5013390 9086065 17272196 17272200 99322575 99322649
1064220 1064225 1064229 1064255 1064274 1064294 1064310 48479461 79747082 114430192
70960163 70960164
28221292 81632947 81632950 81632952 81632953 81632956 81632957 81632959 81632962 81656916 81656919 90963696 98925936 98925951 101107780 101107795 101107796 101107802 101107810 101107814 101130189 101130197
50332288 68125612 77366932 92913528 92913559 92913632 92913633 92913661
4554630 17804039 96126902
37064962 37069815 37070982 37081967 37082002 37082051 37082209 37084592 42133541 42133554 42133560 42133562 62536048 62536050 62536054 62536055 62536068 62536070 62536072 62536074 84175249
75941906
55501564 55501565 55501587 55501590 55501598 55501602 55501606 55501608 55501669 55501672 55501679 55501686 55501687 55501696 55501704 59586669 59586688 59586692 59586697 59586703 59586706 59586716
1628096 1628097 1628487 1628500 1628824 1628842 1628851 1628978 1630468 1630998 1636655 1674871 1674874 1674891 1674904 1674907 1674920 1674927 1674930 1674935 1674939 1710429 1710493 1710499 1710677 64472494
27267720
142631953

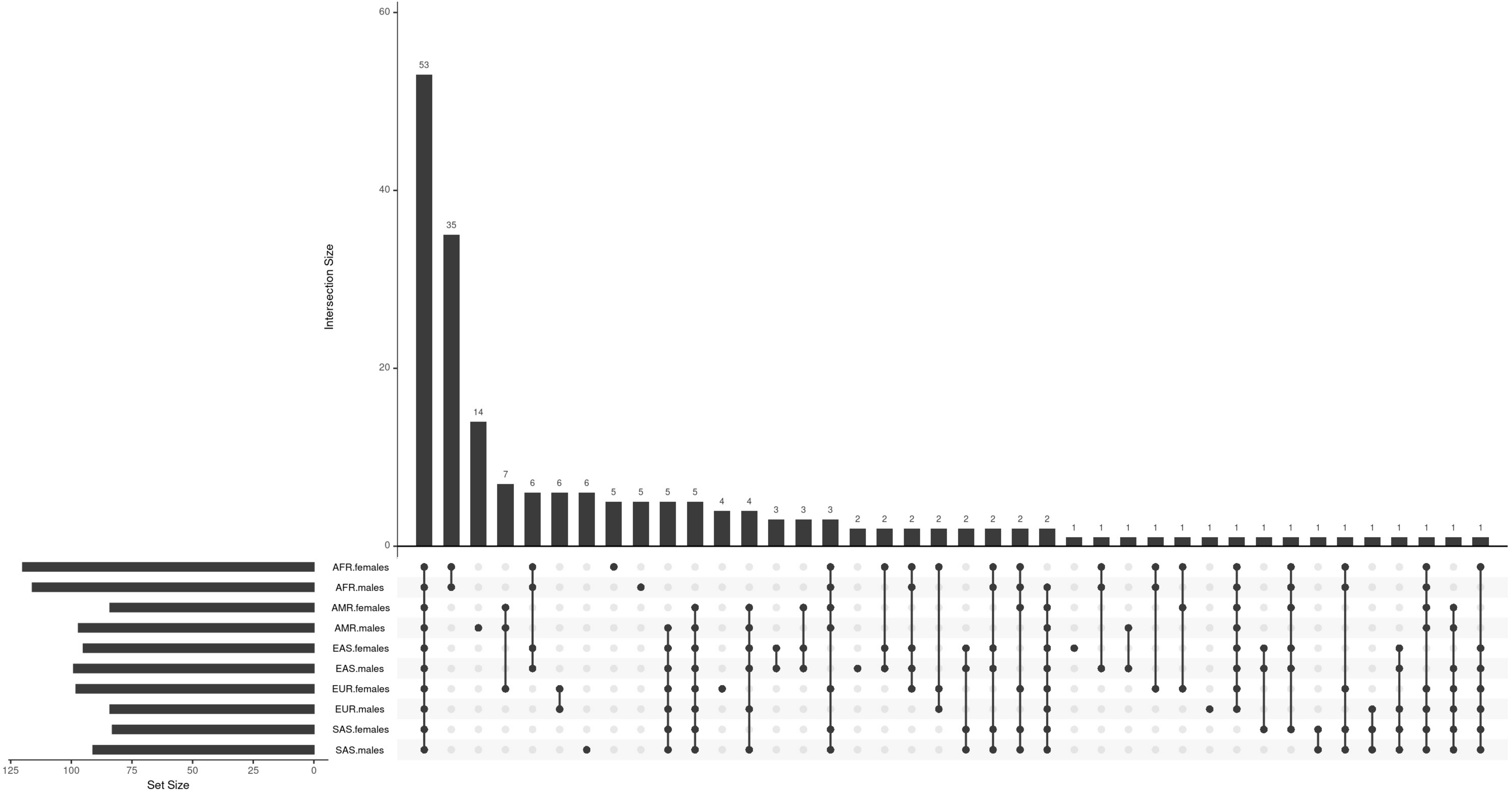
Intersection Size
60
40
20
0
53
35
14
7
6
6
6
5
5
5
5
4
4
3
3
3
2
2
2
2
2
2
2
2
1
1
1
1
1
1
1
1
1
1
1
1
1
1
1
1
AFR.females
AFR.males
AMR.females
AMR.males
EAS.females
EAS.males
EUR.females
EUR.males
SAS.females
SAS.males
125
100
75
50
25
0
Set Size

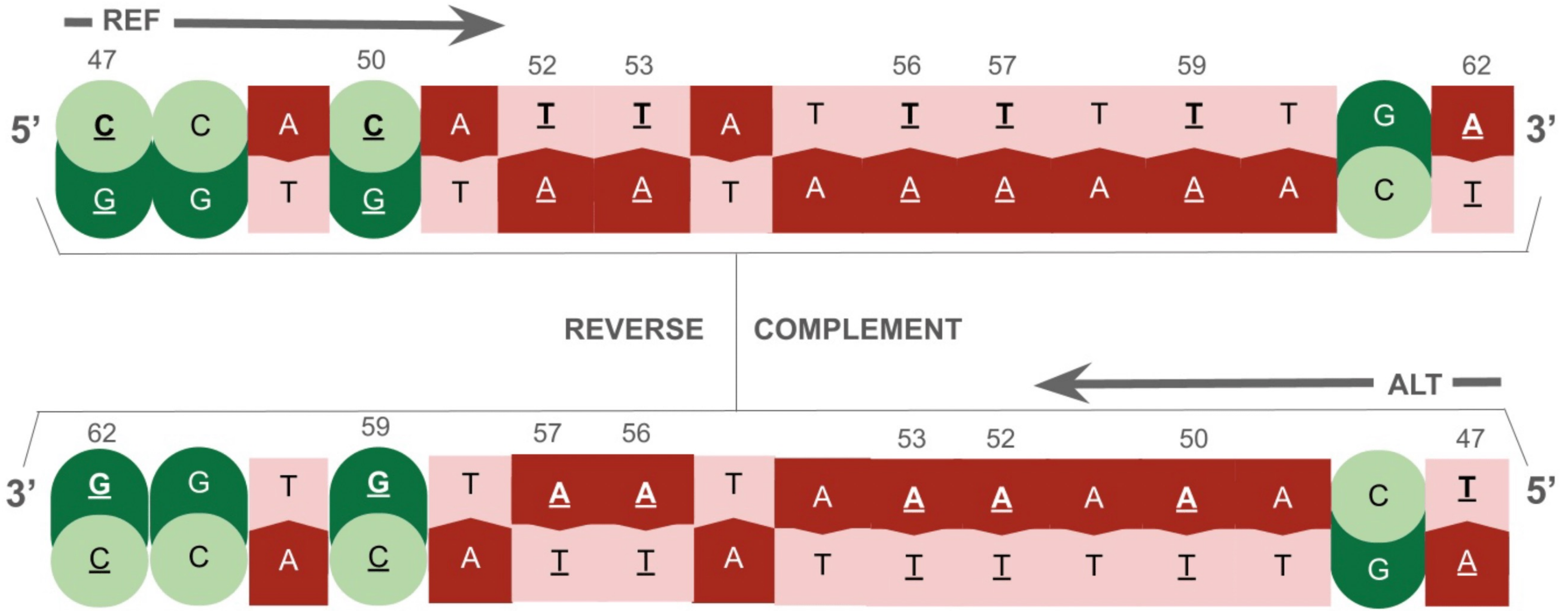
REF
47 50 52 53 56 57 59 62
5' C C A C A T T A T T T T T T G A 3'
G G T G T A A T A A A A A A C T
REVERSE
COMPLEMENT
ALT
62 59 57 56 53 52 50 47
3' G G T G T A A T A A A A A A C T 5'
C C A C A T T A T T T T T T G A

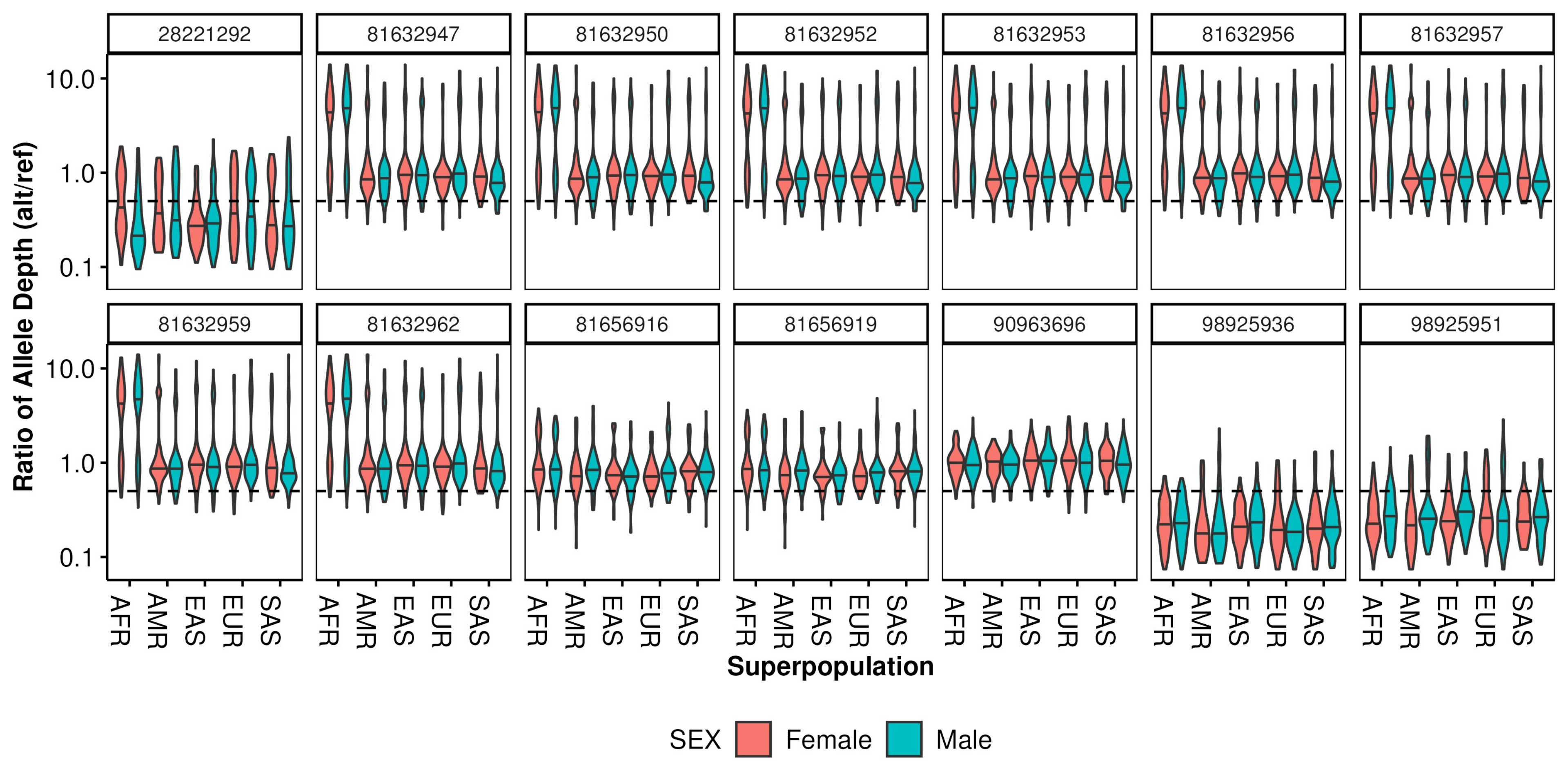
28221292
81632947
81632950
81632952
81632953
81632956
81632957
81632959
81632962
81656916
81656919
90963696
98925936
98925951
Ratio of Allele Depth (alt/ref)
10.0
1.0
0.1
AFR
AMR
EAS
EUR
SAS
Superpopulation
SEX
Female
Male

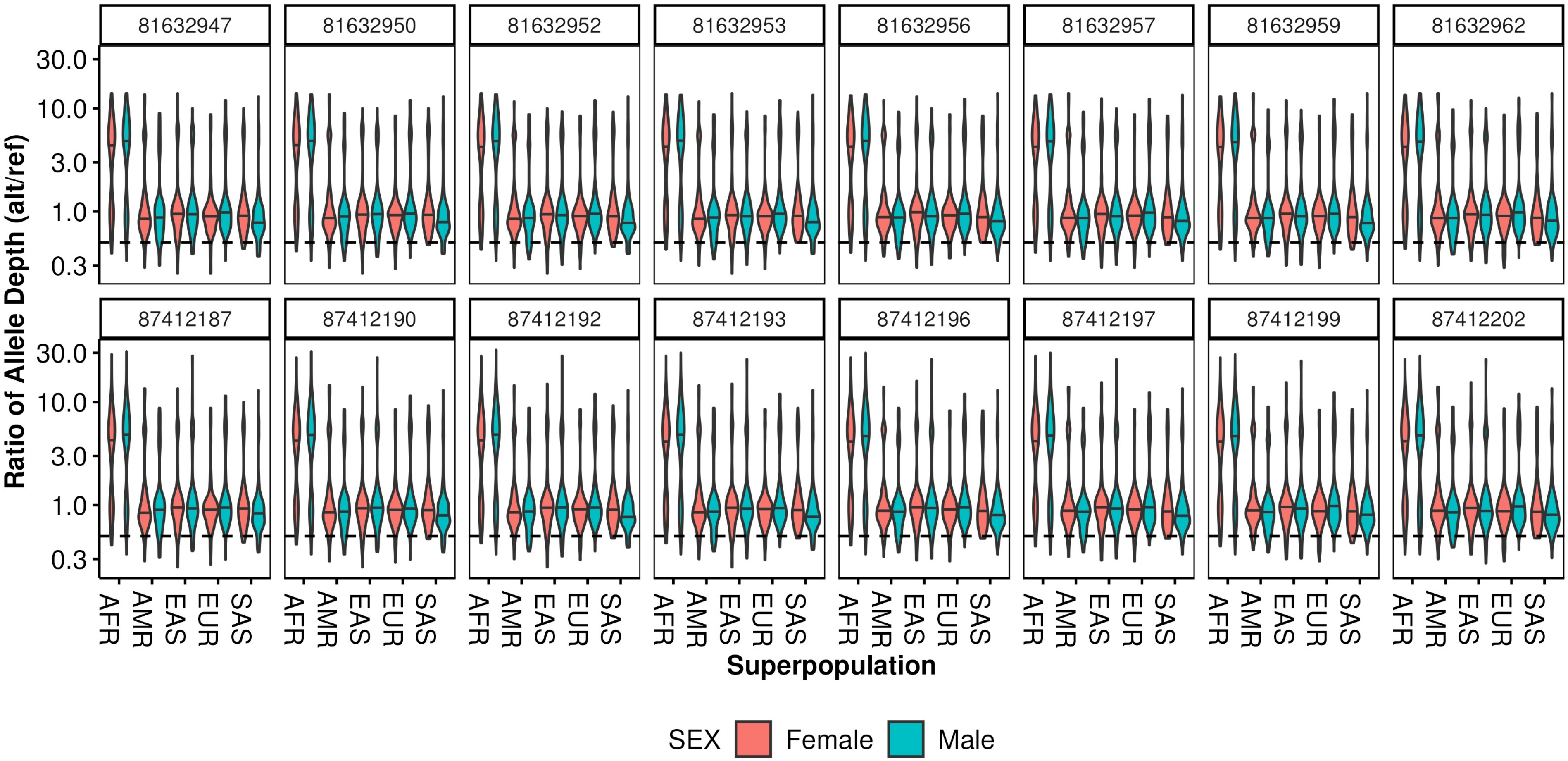

81632947
81632950
81632952
81632953
81632956
81632957
81632959
81632962
87412187
87412190
87412192
87412193
87412196
87412197
87412199
87412202
30.0
10.0
3.0
1.0
0.3
Ratio of Allele Depth (alt/ref)
AFR
AMR
EAS
EUR
SAS
Superpopulation
SEX
Female
Male

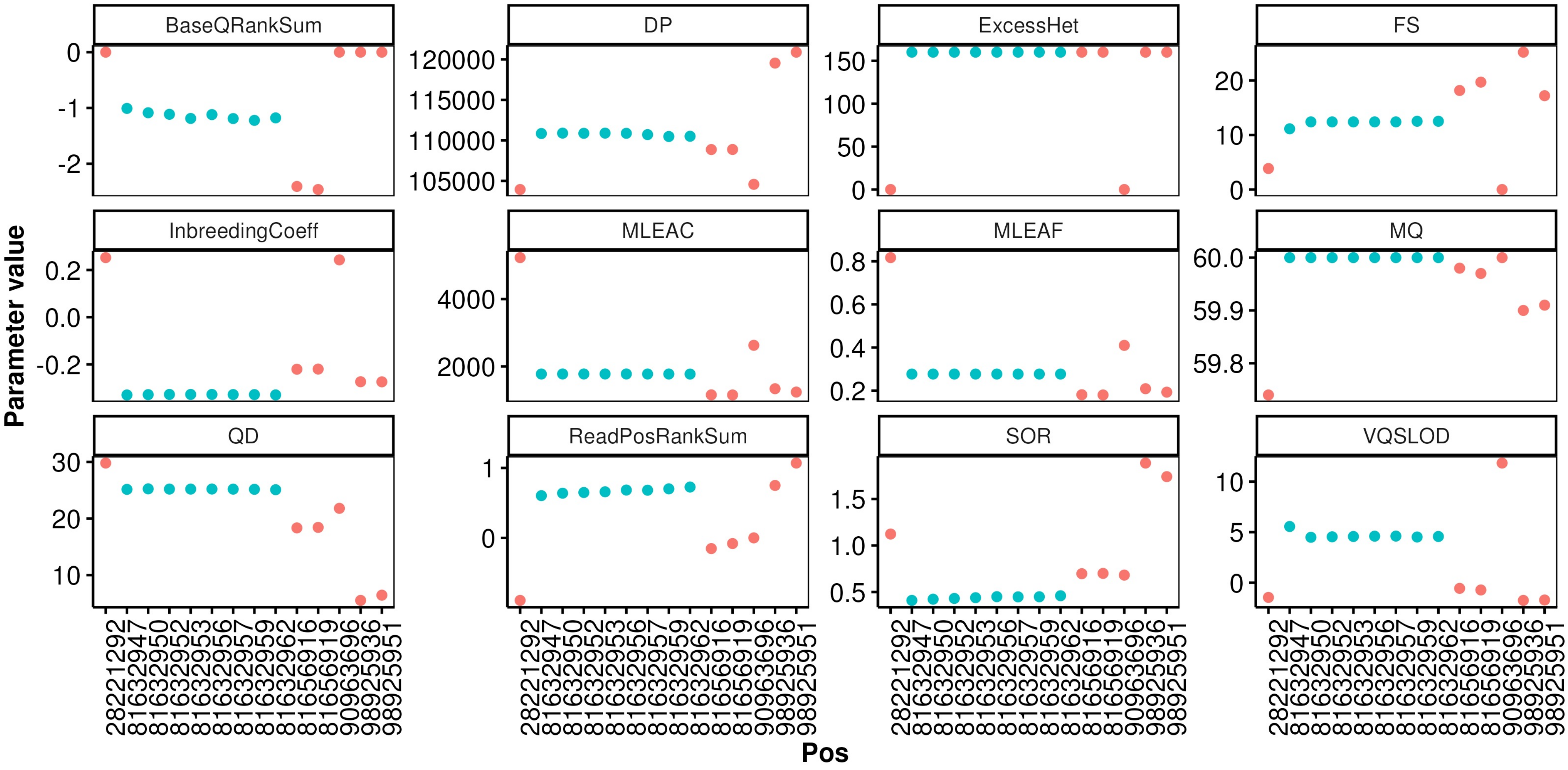

BaseQRankSum
DP
ExcessHet
FS
InbreedingCoeff
MLEAC
MLEAF
MQ
QD
ReadPosRankSum
SOR
VQSLOD
Parameter value
Pos
28221292
81632947
81632950
81632952
81632953
81632956
81632957
81632959
81632962
81656916
81656919
90963696
98925936
98925951

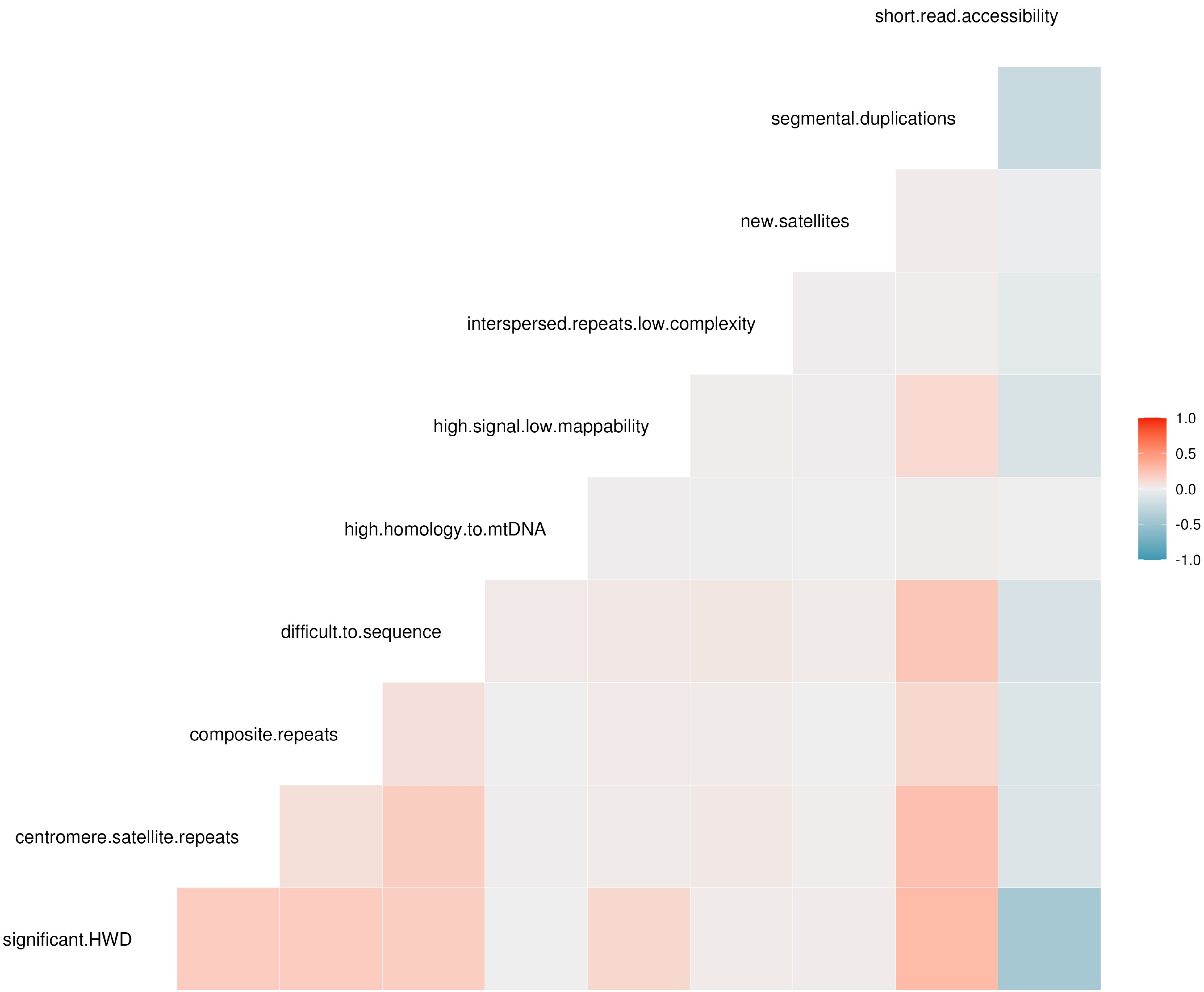
short.read.accessibility
segmental.duplications
new.satellites
interspersed.repeats.low.complexity
high.signal.low.mappability
high.homology.to.mtDNA
difficult.to.sequence
composite.repeats
centromere.satellite.repeats
significant.HWD
1.0
0.5
0.0
-0.5
-1.0

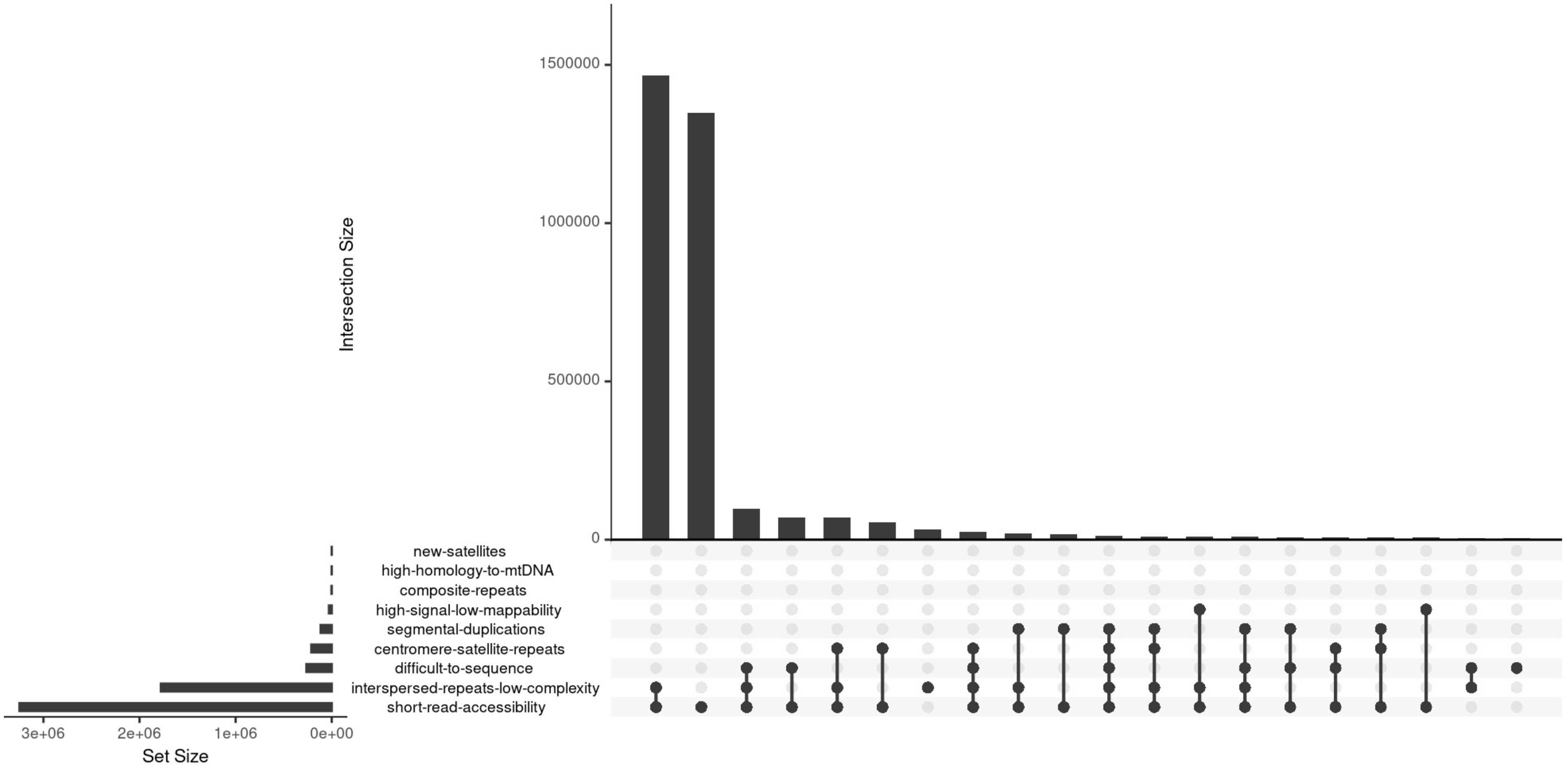

Intersection Size
1500000
1000000
500000
0
new-satellites
high-homology-to-mtDNA
composite-repeats
high-signal-low-mappability
segmental-duplications
centromere-satellite-repeats
difficult-to-sequence
interspersed-repeats-low-complexity
short-read-accessibility
3e+06
2e+06
1e+06
0e+00
Set Size

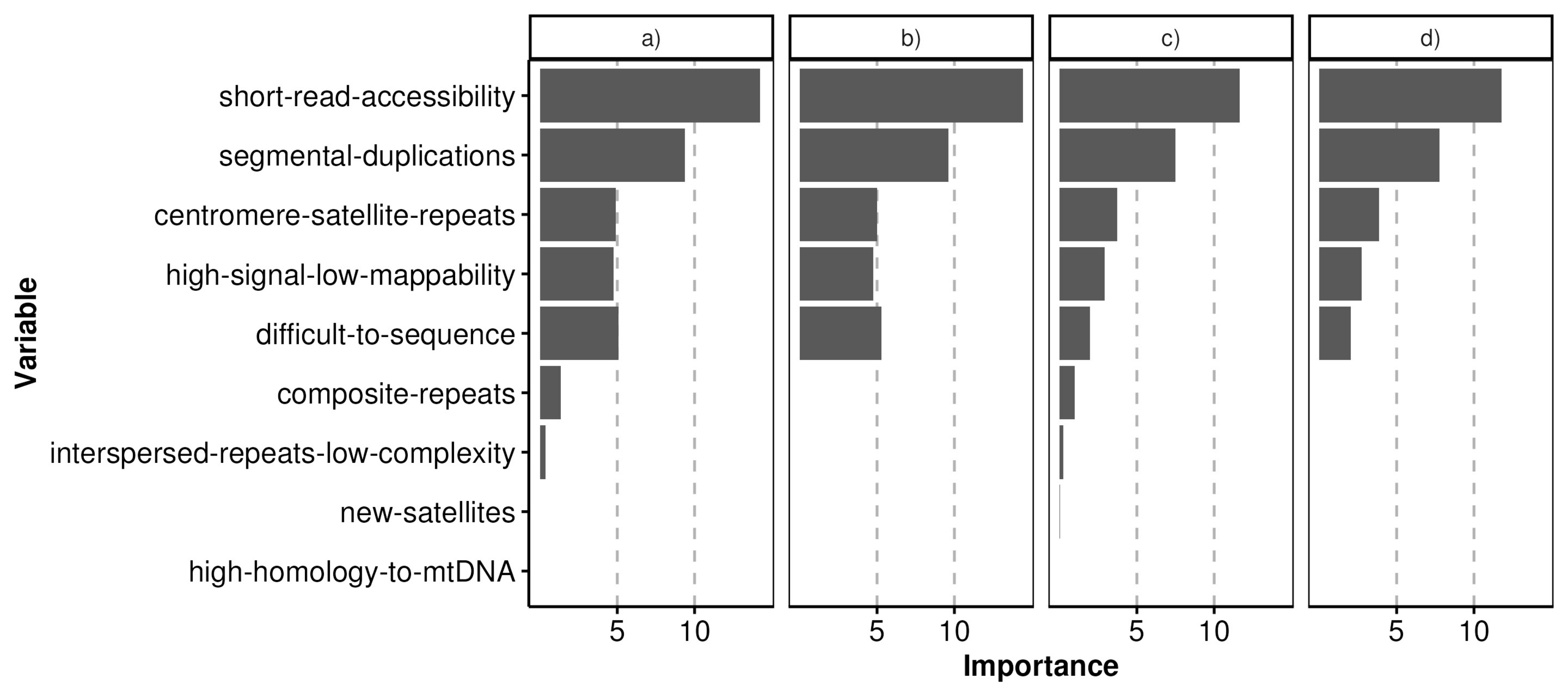

a)
b)
c)
d)
short-read-accessibility
segmental-duplications
centromere-satellite-repeats
high-signal-low-mappability
difficult-to-sequence
composite-repeats
interspersed-repeats-low-complexity
new-satellites
high-homology-to-mtDNA
Variable
5
10
Importance

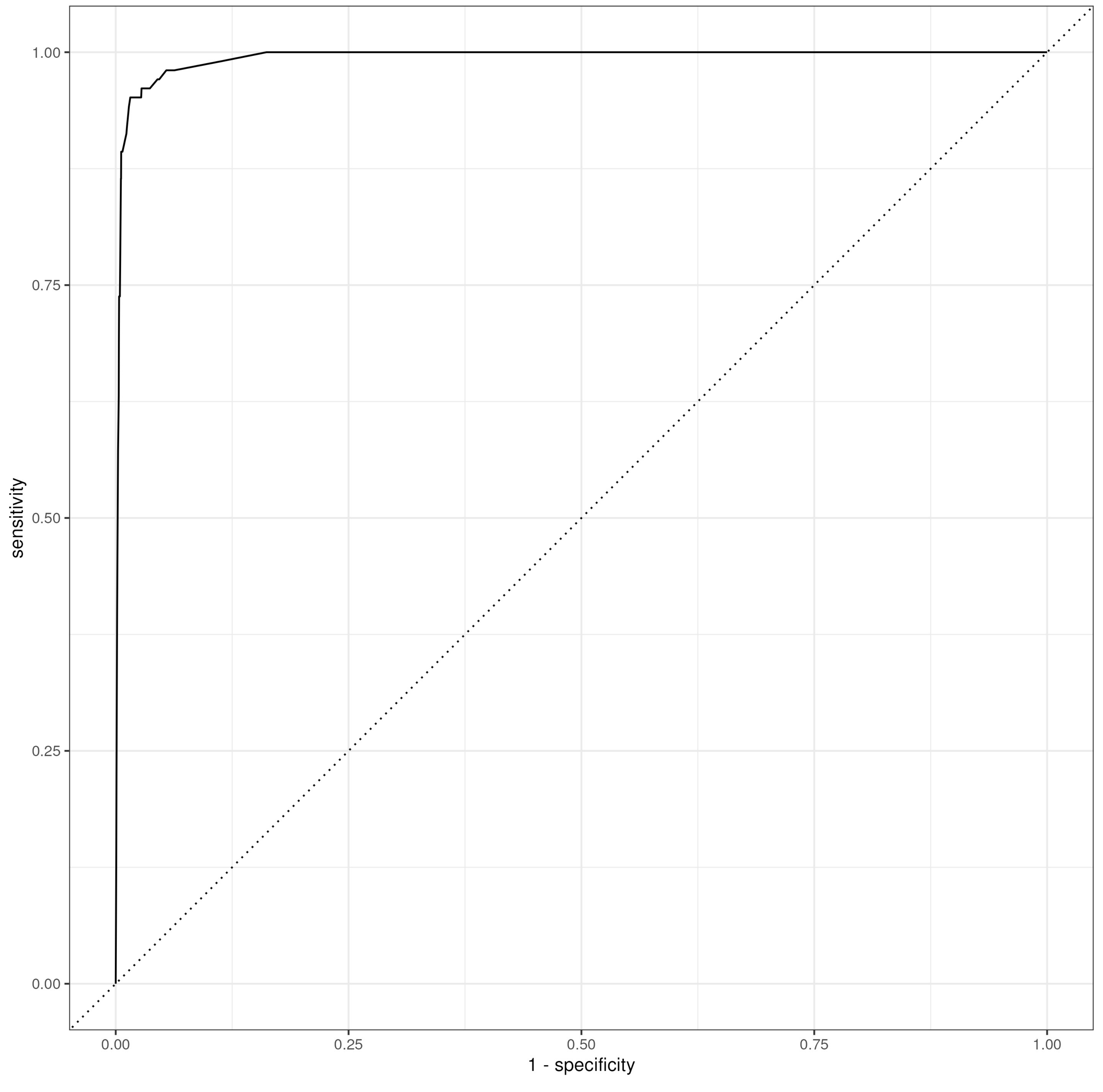
1.00
0.75
0.50
0.25
0.00
sensitivity
0.00
0.25
0.50
0.75
1.00
1 - specificity

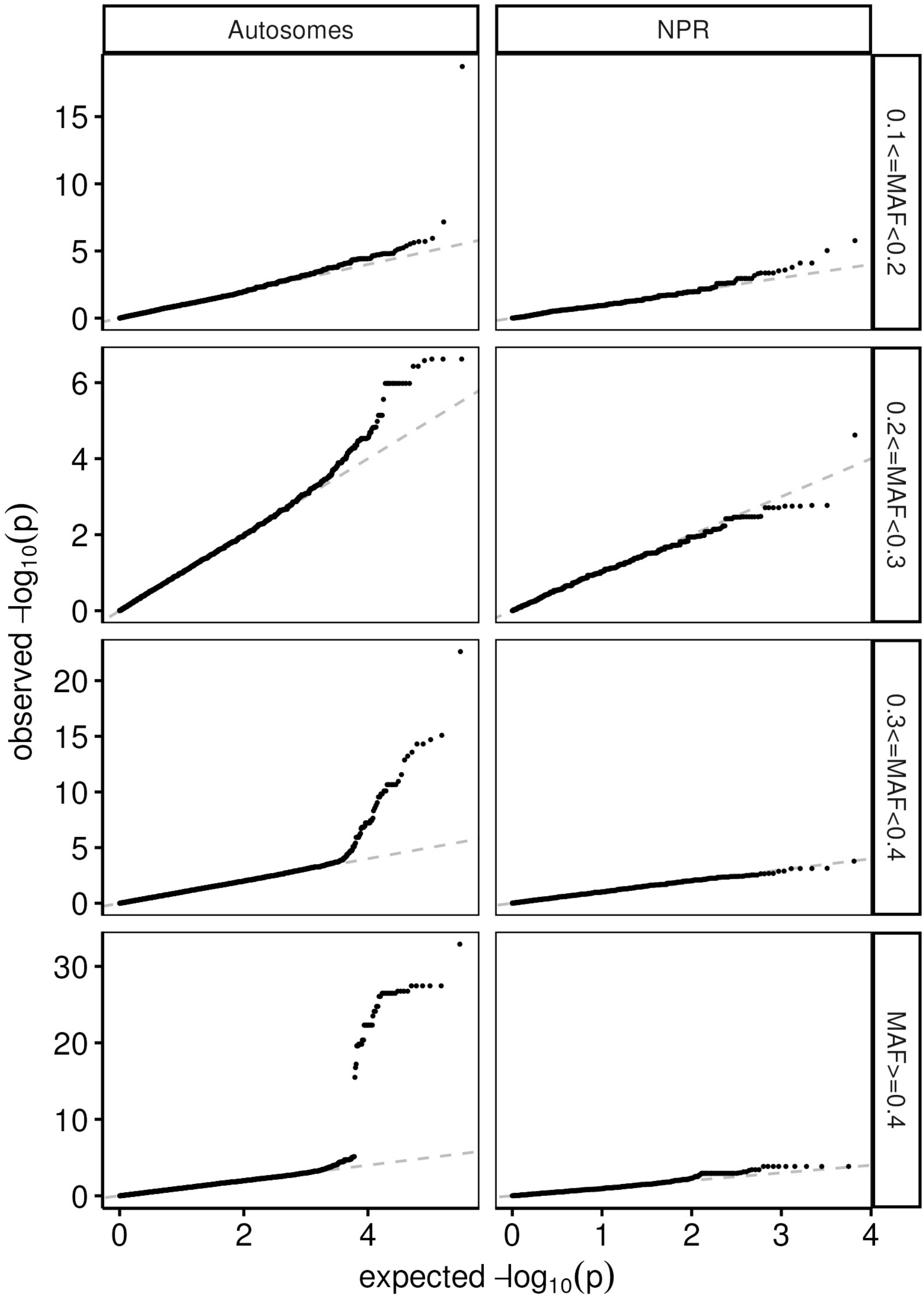

Autosomes
NPR
0.1<=MAF<0.2
0.2<=MAF<0.3
0.3<=MAF<0.4
MAF>=0.4
observed -log10(p)
expected -log10(p)

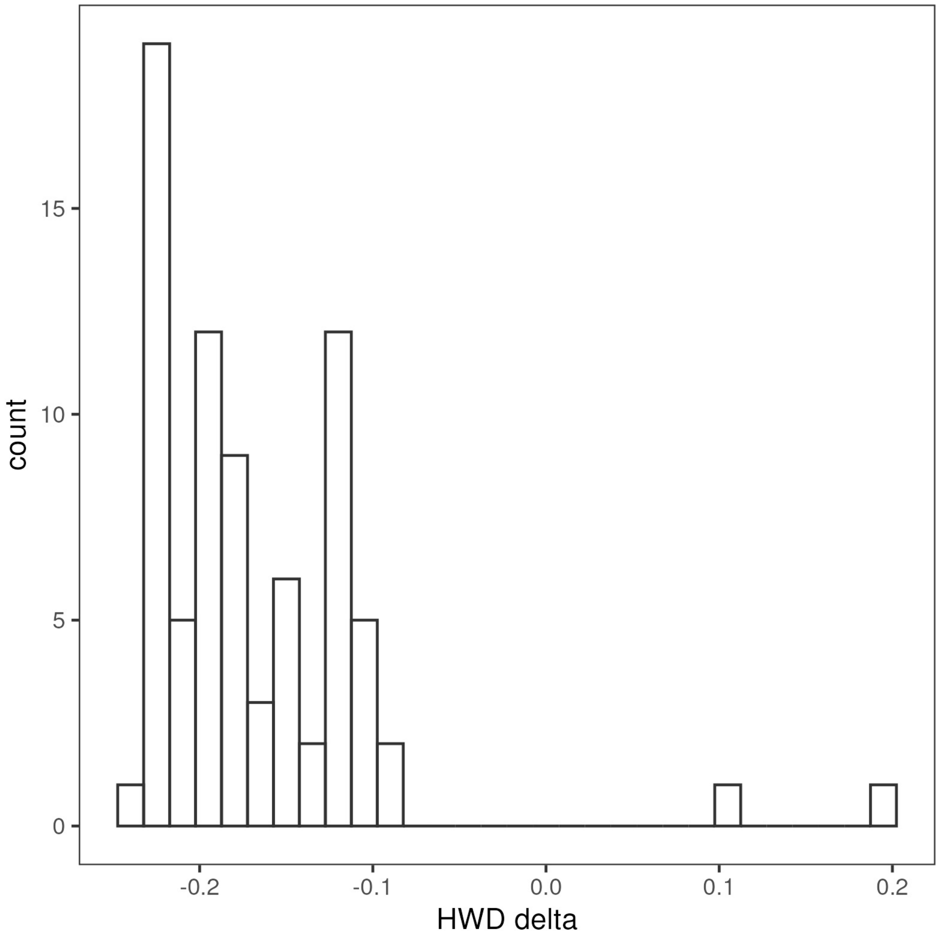
count
15
10
5
0
-0.2
-0.1
0.0
0.1
0.2
HWD delta

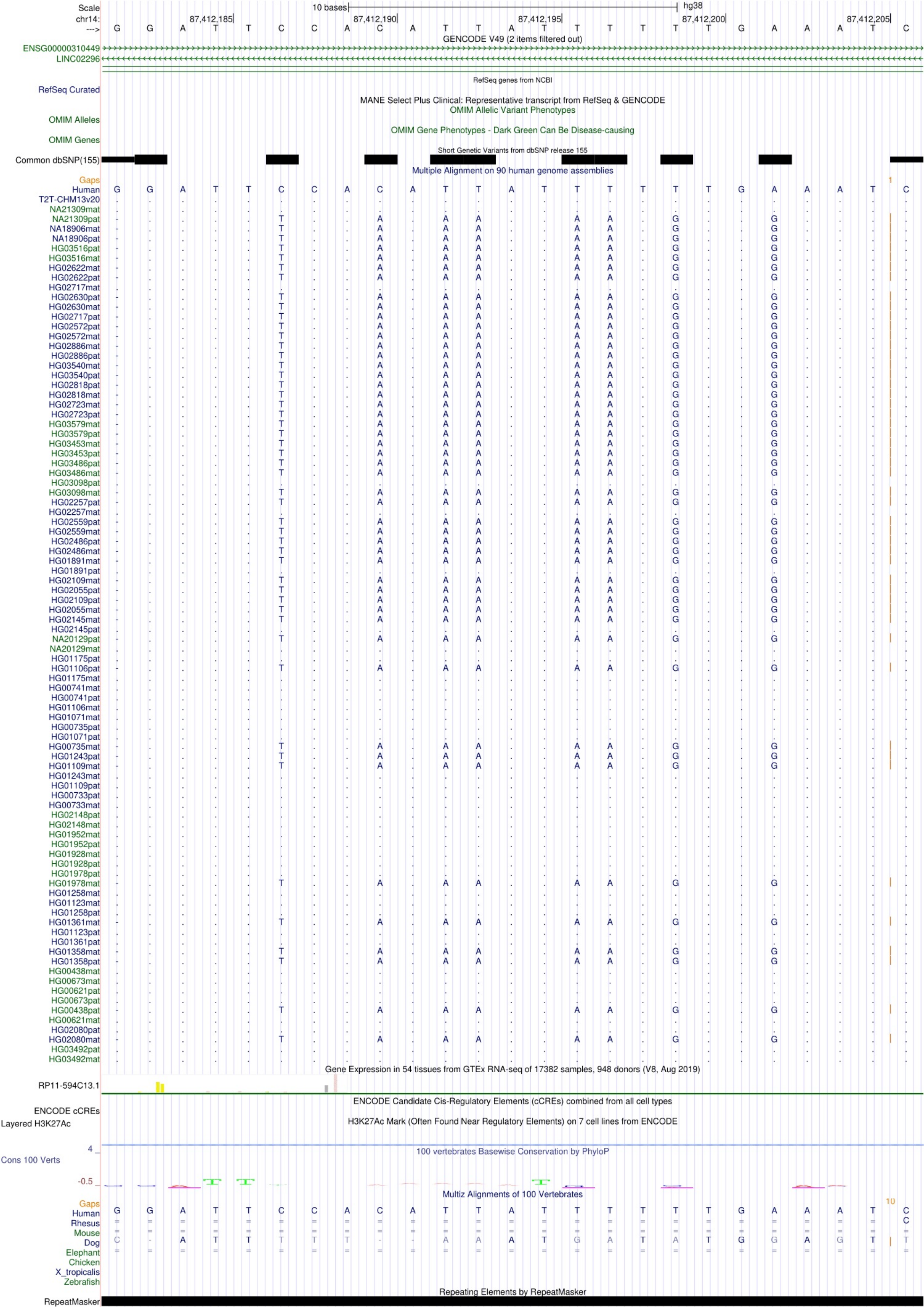

Scale
10 bases
hg38
chr14:
87,412,185
87,412,190
87,412,195
87,412,200
87,412,205
--->
G G A T T C C A C A T T A T T T T T T G A A A T C
GENCODE V49 (2 items filtered out)
ENSG00000310449
LINC02296
RefSeq genes from NCBI
RefSeq Curated
MANE Select Plus Clinical: Representative transcript from RefSeq & GENCODE
OMIM Allelic Variant Phenotypes
OMIM Alleles
OMIM Gene Phenotypes - Dark Green Can Be Disease-causing
OMIM Genes
Short Genetic Variants from dbSNP release 155
Common dbSNP(155)
Multiple Alignment on 90 human genome assemblies
Gaps
Human
T2T-CHM13v20
NA21309mat
NA21309pat
NA18906mat
NA18906pat
HG03516pat
HG03516mat
HG02622mat
HG02622pat
HG02717mat
HG02630pat
HG02630mat
HG02717pat
HG02572pat
HG02572mat
HG02886mat
HG02886pat
HG03540mat
HG03540pat
HG02818pat
HG02818mat
HG02723mat
HG02723pat
HG03579mat
HG03579pat
HG03453mat
HG03453pat
HG03486pat
HG03486mat
HG03098pat
HG03098mat
HG02257pat
HG02257mat
HG02559pat
HG02559mat
HG02486pat
HG02486mat
HG01891mat
HG01891pat
HG02109mat
HG02055pat
HG02109pat
HG02055mat
HG02145mat
HG02145pat
NA20129pat
NA20129mat
HG01175pat
HG01106pat
HG01175mat
HG00741mat
HG00741pat
HG01106mat
HG01071mat
HG00735pat
HG01071pat
HG00735mat
HG01243pat
HG01109mat
HG01243mat
HG01109pat
HG00733pat
HG00733mat
HG02148pat
HG02148mat
HG01952mat
HG01952pat
HG01928mat
HG01928pat
HG01978pat
HG01978mat
HG01258mat
HG01123mat
HG01258pat
HG01361mat
HG01123pat
HG01361pat
HG01358mat
HG01358pat
HG00438mat
HG00673mat
HG00621pat
HG00673pat
HG00438pat
HG00621mat
HG02080pat
HG02080mat
HG03492pat
HG03492mat
Gene Expression in 54 tissues from GTEx RNA-seq of 17382 samples, 948 donors (V8, Aug 2019)
RP11-594C13.1
ENCODE Candidate Cis-Regulatory Elements (cCREs) combined from all cell types
ENCODE cCREs
H3K27Ac Mark (Often Found Near Regulatory Elements) on 7 cell lines from ENCODE
Layered H3K27Ac
4
100 vertebrates Basewise Conservation by PhyloP
Cons 100 Verts
-0.5
Multiz Alignments of 100 Vertebrates
Gaps
10
Human
Rhesus
Mouse
Dog
Elephant
Chicken
X_tropicalis
Zebrafish
Repeating Elements by RepeatMasker
RepeatMasker

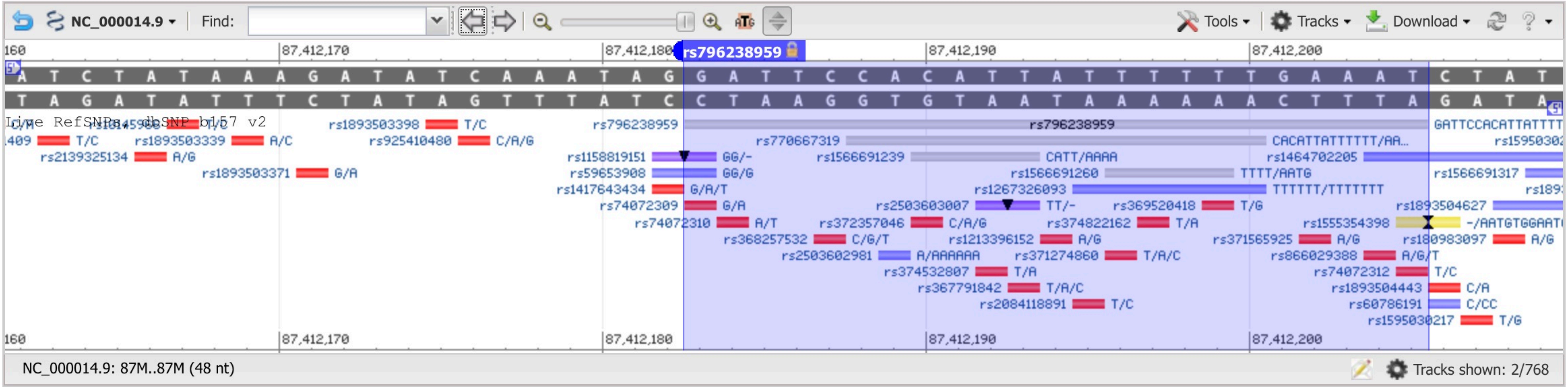

NC_000014.9
Find:
Tools
Tracks
Download
87,412,170
87,412,180
rs796238959
87,412,190
87,412,200
A T C T A T A A A G A T A T C A A A T A G G A T T C C A C A T T A T T T T T T G A A A T C T A T
T A G A T A T T T C T A T A G T T T A T C C T A A G G T G T A A T A A A A A A C T T T A G A T A
Live RefSNPs, dbSNP b157 v2
rs1893503398 T/C
rs1893503339 A/C
rs925410480 C/A/G
rs2139325134 A/G
rs1893503371 G/A
rs796238959
rs770667319
rs1158819151 GG/-
rs1566691239 CATT/AAAA
rs59653908 GG/G
rs1566691260 TTTT/AATG
rs1417643434 G/A/T
rs1267326093 TTTTTT/TTTTTTT
rs74072309 G/A
rs2503603007 TT/-
rs369520418 T/G
rs74072310 A/T
rs372357046 C/A/G
rs374822162 T/A
rs368257532 C/G/T
rs1213396152 A/G
rs2503602981 A/AAAAAA
rs371274860 T/A/C
rs374532807 T/A
rs367791842 T/A/C
rs2084118891 T/C
rs1464702205
rs1566691317
rs1893504627
rs1555354398 -/AATGTGGAAT
rs371565925 A/G
rs180983097 A/G
rs866029388 A/G/T
rs74072312 T/C
rs1893504443 C/A
rs60786191 C/CC
rs1595030217 T/G
NC_000014.9: 87M..87M (48 nt)
Tracks shown: 2/768

```
rs796238959       CACATTATTTTTT
                  |||||||||||||
rs770667319  GATTCCACATTATTTTTTGAAAT
```
