## Supplementary Captions for "Assessing Hardy-Weinberg Equilibrium in T2T-aligned 1000 Genomes Project"

### Supporting Information captions

#### Supplementary figures

**Figure S1. QQ plots and p-value histograms for autosome-wide HWE in each of the 10 groups.** HWE results from five super-populations and two sexes are shown for autosomal SNPs with MAF>=5% (GC $\lambda$ for females: AFR = 1.12, AMR = 1.28, EAS = 1.11, EUR = 1.08, SAS = 1.17; males: AFR = 1.10, AMR = 1.32, EAS = 1.10, EUR = 1.10, SAS = 1.14).

**Figure S2. QQ plots and p-value histograms for autosome-wide HWE meta-analysis of 10 groups.** Omnibus meta-analysis results from five super-populations and two sexes are shown for autosomal SNPs which have MAF>=5% in all groups and fall in: a) all regions (GC $\lambda$ = 1.38) b) reliable regions (GC $\lambda$ = 1.33).

**Figure S3. QQ plots and p-value histograms for X-chromosome HWE in each of the 10 groups.** HWE results from five super-populations and two sexes are shown for chrX SNPs with MAF>=5% (GC $\lambda$ for females: AFR = 1.19, AMR = 1.56, EAS = 1.11, EUR = 1.12, SAS = 1.27; males: AFR = 1.11, AMR = 1.36, EAS = 1.18, EUR = 1.00, SAS = 1.09). NPR data is only for females.

**Figure S4. QQ plots and p-value histograms for X-chromosome HWE meta-analysis of 10 groups.** Omnibus meta-analysis results from five super-populations and two sexes are shown for chrX SNPs which have MAF>=5% in all groups and fall in: a) all regions (GC $\lambda$ = 1.55) b) reliable regions (GC $\lambda$ = 1.44). NPR data is only for females. Genomic features were not available for PARs except difficult-to-sequence regions.

**Figure S5. Heatmap of HWD delta for 256 SNPs with significant HWD.** HWD delta is shown by a signed color scale for each of the 10 groups at the 256 SNPs (Table S7). Panels represent chromosomes and genomic location with each chromosome is arranged in ascending order on the X-axis. Red color represents negative delta, which means excess of heterozygotes. See companion Figure 6 for gradient colors.

**Figure S6. Upset plot for 256 SNPs with significant HWD.** HWD significance in each of the 10 groups is compared across the 256 SNPs (Table S7). Sets are arranged alphabetically and intersections are arranged by frequency.

**Figure S7. Reverse complement sequence at chr14 locus.** The reference and alternate sequences at the chr14:81,632,947-81,632,962 position are reverse complements of each other. SNPs that are highlighted with the last two digits of the base pair positions are the 8 SNPs that were observed to have HWD in our analysis.

**Figure S8. Allele depth at chr14 locus in T2T compared to neighbouring HWD SNPs.** Ratio of alternative and reference allele depth is shown for the chr14 locus (chr14:81,632,947-81,632,962) and neighbouring SNPs with HWD (locations in Table S7) in the 10 groups. Dashed line indicates a ratio of 0.5. Y-axis is presented on a log10 scale.

**Figure S9. Allele depth at chr14 locus in T2T compared to GRCh38.** Ratio of alternative and reference allele depth is shown for the chr14 locus (chr14:81,632,947-81,632,962) on T2T (top) with corresponding location on GRCh38 (bottom) in the 10 groups. Dashed line indicates a ratio of 0.5. Y-axis is presented on a log10 scale.

**Figure S10. Quality parameters of chr14 locus in T2T compared to neighbouring HWD SNPs.** Site-specific parameters are shown for the chr14 locus (chr14:81,632,947-81,632,962 in blue) and neighbouring SNPs with HWD (locations in Table S7).

**Figure S11. Correlation between autosome-wide genomic features.** Pairwise Spearman correlation between genomic features in the autosome is shown by a signed color scale. Correlation between each feature and significant HWD status is also shown in the bottom row.

**Figure S12. Upset plot for autosome-wide genomic features.** Overlap of genomic feature positions is examined across analyzed autosomal SNPs (n=3,377,201). They map on to 3,325,907 unique positions; 51,294 positions are not covered by any of the genomic features examined. Sets are arranged alphabetically and intersections are arranged by frequency; top 20 intersections are shown.

**Figure S13. Variable importance of genomic features in the predictive autosome-wide model.** Variable importance is shown for: a) 9 features in training set, b) 5 selected features in training set, c) 9 features in test set, d) 5 selected features in test set.

**Figure S14. ROC curve for the predictive autosome-wide model of genomic features.** A multivariate logistic model to predict binary HWD significance status was trained on odd autosomes and tested on even autosomes.

**Figure S15. QQ plots for autosomes and NPR HWE of recently-admixed groups.** HWE results from two populations (recently-admixed groups ASW and ACB) are shown for autosomal and chrX SNPs with MAF>=10%. NPR data is only for females. SNPs are stratified by MAF (GC $\lambda$ for autosomes, chrX): 0.1<=MAF<0.2 (1.02,1.09), 0.2<=MAF<0.3 (1.07,1.11), 0.3<=MAF<0.4 (1.01,1.18), MAF>=0.4 (1.00,0.89).

**Figure S16. Histogram of HWD delta of significant SNPs in recently-admixed groups.** HWE results from two populations (recently-admixed groups ASW and ACB) are shown for autosomal SNPs with MAF>=10% and HWE p-value < 5e-8 (Table S10).

**Figure S17. UCSC genome browser at chr14 locus.** Pangenome variation at the chr14 locus is examined in the UCSC genome browser (link in Resources).

**Figure S18. Variation viewer at chr14 locus.** A high density of variation is seen at the chr14 locus in dbSNP (build 157) Variation Viewer.

**Figure S19. MNVs at chr14 locus.** Alignment of the two MNVs (rs796238959 and rs770667319) at chr14 locus.

#### Supplementary Tables

**Table S1. SNPs with significant HWE heterogeneity but not-significant omnibus test results.** Omnibus, HWE heterogeneity and sdMAF results are shown for 7 SNPs (from Table 2) along with group-specific details, and a column to indicate if they fall in reliable regions.

**Table S2. HWD in autosome-wide genomic features. I**mpact of each genomic feature on autosomal HWD is shown by chi-squared test and odds-ratio. Two genomic features (“assembly gaps” and “telomere”) had no overlaps with the analyzed genome and hence are not displayed here.

**Table S3. Genomic details of genome-wide SNPs with HWD in reliable regions.** 255 SNPs on autosomes and 1 on NPR with significant HWD were found in reliable regions. Their coordinates were lifted over to GRCh38 for functional annotations from dbSNP and Ensembl VEP (see Methods). An additional annotation indicates their QC status from the original work (2).

**Table S4. GWAS catalog results of genome-wide SNPs with HWD in reliable regions.** 10 SNPs on autosomes had a significant result in the NHGRI-EBI GWAS catalog with $p<5e-8$.

**Table S5. Summary of GWAS catalog results of genome-wide SNPs with HWD in reliable regions.** The number of studies associated with each of the 10 SNPs with $p<5e-8$ in the NHGRI-EBI GWAS catalog.

**Table S6. Group-wise details of genome-wide SNPs with HWD in reliable regions.** Allele counts, MAF and HWE results are shown for the 256 SNPs in each of the 10 groups.

**Table S7. Meta-analysis results of genome-wide SNPs with HWD in reliable regions.** Omnibus, HWE heterogeneity and sdMAF results are shown for the 256 SNPs along with group-specific details from Table S6.

**Table S8. Meta-analysis and group-wise details results of 1 SNP with sdHWD in PAR2.** Omnibus, HWE heterogeneity and sdMAF results are shown for 1 PAR2 SNP along with sex-specific details for the EUR super-population group where it was found.

**Table S9. Results for the multivariate genomic feature model for autosomal HWD.** A multivariate logistic model to predict binary HWD significance status was trained on odd autosomes and tested on even autosomes.

**Table S10. HWE results of significant SNPs in recently-admixed populations.** HWE results from two populations (recently-admixed groups ASW and ACB) are shown for autosomal SNPs with MAF>=10% and HWE p-value < 5e-8 (Figure S13).

**Table S11. Sources for the genomic features used in analysis.** Resource table for genomic features with links to the resources. Last column indicates if that genomic feature was available for PARs.
